# Multiscale patterning of a model apical extracellular matrix revealed by systematic endogenous protein tagging

**DOI:** 10.1101/2025.05.14.653803

**Authors:** James Matthew Ragle, Murugesan Pooranachithra, Guinevere E. Ashley, Emma Cadena, Batia Blank, Katelyn Kang, Cincy Chen, Aditree R. Bhowmick, Soraya H. Mercado, Tabatha E. Wells, John C. Clancy, Andrew D. Chisholm, Jordan D. Ward

**Affiliations:** Department of Molecular Cell and Developmental Biology, University of California Santa Cruz, Santa Cruz, CA 95064; Department of Cell and Developmental Biology, School of Biological Sciences, University of California San Diego, La Jolla, CA 92093

**Keywords:** collagen, ZP protein, ecdysis, matrisome, CRISPR

## Abstract

Barrier epithelia are shielded from the external environment by their apical extracellular matrices (aECMs). The molecular complexity of aECMs has challenged understanding of their organization *in vivo*. To define the molecular architecture of a model aECM we generated a toolkit of 102 fluorescently tagged aECM components using gene editing in *C. elegans*, focusing on proteins secreted by the epidermis to form the collagen-rich cuticle. We developed efficient pipelines for modular protein tagging and rapid fluorophore swapping. Most tagged collagens were functional and exhibited exquisitely specific patterning across stages, cell types, and matrix substructures. We define multiple reference markers for key substructures including the little-understood cortical layer, as well as the helical crossed fiber arrays that function as a hydrostatic skeleton to maintain organismal shape. We further tagged >30 members of key aECM protein classes including proteases, protease inhibitors, and lipid transporters. Our standardized markers will allow dissection of the mechanistic basis of aECM spatiotemporal patterning *in vivo*.

**Highlights:**

- First large-scale protein tagging resource for the apical extracellular matrix
- Optimization of CRISPR methods for protein tagging including color swaps
- Tagged proteins are functional and exhibit a high degree of stage-, cell- and compartment specificity
- Reference localization patterns for multiple aECM compartments and markers for newly defined compartments

## Introduction

Extracellular matrix (ECM) comprises a wide variety of compartments including those formed by epithelial tissues such as basement membranes (BMs) and apical extracellular matrices (aECMs). ECMs play myriad roles in organismal development,^5^ and are key determinants of disease pathology^8^ and aging.^10^ The molecular complexity and diversity of such ECMs is increasingly appreciated. Estimates of the total number of protein species of the ECM, i.e. the matrisome,^11,12^ range from the hundreds to over 1000, many of which are subject to complex processing and post-translational modification. In short, the ECM is one of the most complex aspects of metazoan development and physiology.

One major type of ECM is the basement membrane of epithelia, formed around a conserved set of proteins such as laminins and type IV collagens.^13^ Recent studies in *C. elegans* and mammals have systematically tagged 46 BM components including tissue-or stage-specific proteins.^14,15^ These large-scale resources have revealed the molecular diversity of BMs *in vivo*. While previous work had suggested that BMs turned over on a scale of weeks, analysis of endogenously tagged proteins revealed BMs are far more dynamic than appreciated.^14^ Such resources have also established the importance of studying ECM components using endogenous knock-in tags, consistent fluorescent protein (XFP) tags and a modular editing pipeline for stable and reproducible analysis of protein localization.

Here we focus on apical extracellular matrices (aECMs), a highly heterogeneous set of complex ECMs.^16^ The nematode *C. elegans* generates several distinct aECMs,^17,18^ of which the largest is the epidermal aECM or cuticle, a critical determinant of organismal morphogenesis,^19^ permeability barrier function,^20^ skin wound healing,^21^ mechanosensation,^22^ host-pathogen interactions,^23,24^ mate recognition,^25^ and aging.^10^ The main body cuticle is continuous with more specialized interfacial cuticles.^26^ Throughout development the cuticle can be broadly divided into collagen-rich basal layers and cortical layers containing collagens and additional insoluble cuticlins and epicuticlins.^17^ The surface of the cortical layer further forms a lipid-rich epicuticle that contributes to the permeability barrier^27^ and an outermost glycoprotein-rich surface coat that affects interaction with pathogens.^28^ The main body cuticle is patterned with circumferential annuli separated by furrows. The body cuticle also contains multiple stage- and region-specific specializations, such as the longitudinal ridges (alae) formed in L1, dauer, and adult^29^. Compared to the transient cuticles formed in development, the mature adult cuticle contains several additional compartments such as a fluid-filled medial layer between the cortical and fibrous layers that contains organized arrays of cylindrical struts.^7^

The *C. elegans* epidermal aECM is remarkably complex at a molecular level, with at least 175 distinct types of cuticle collagens and a similar number of non-collagen proteins ^9^ such as cuticlins, epicuticlins, C-type lectins, chondroitin proteoglycans, mucins, and zona pellucida (ZP) domain proteins. Specific collagen localization correlates with known morphological sub-structures such as furrows or annuli.^30^ Transcriptomic studies have revealed that ∼20% of *C. elegans* genes oscillate in expression, peaking at a stereotypical point in each larval stage.^3,31,32^ Such gene sets are enriched in aECM components whose oscillation reflects the periodic assembly of the cuticle in the four larval molts. Mutation or inactivation of specific oscillating collagens affects the structural organization of collagens expressed at a similar time, but not collagens from other temporal groups.^30^ Moreover, the epidermal aECM of each stage is prefigured by a transient aECM known as precuticle.^17^ Super-resolution microscopy has revealed nanoscale organization of collagens within the adult-specific strut compartment^7^ and has revealed epicuticlins as novel components of the cortical layer.^21^ These studies have reinforced the importance of standardized tagging to build a molecular atlas of aECM components.

Here we report a protein-tagging resource of knock-in strains for 102 aECM components as the basis for such a molecular atlas. We selected genes likely to be informative aECM markers based on several criteria including genetic function, expression pattern, or sequence orthology. We optimized a gene editing pipeline to make endogenously mNeonGreen (mNG) tagged versions of 65 cuticle collagens and 25 non-collagen aECM proteins. For select genes, we generated an additional mScarlet (mSc) red FP-tagged versions to facilitate co-localization studies. We tagged 40 proteins previously shown to be components of cuticle and validated previously observed patterns. Of the set of collagens identified based on phenotype, most tagged knock-ins retained normal function. While many tagged proteins localized to previously known cuticle substructures, some defined previously unknown substructures or revealed previously unappreciated aspects of known structures. The aECM toolkit provides a resource for further studies of aECM architecture and the mechanistic basis of complex ECM structure and function.

## Results

### A modular pipeline for protein-tagging knock-inss

We first optimized parameters for Cas9 ribonucleoprotein complex (RNP) and linear repair template-based genome editing using a GFP knock-in into the 5’ end of the *glh-1* gene, similar to a previous optimization study.^1^ We tested Cas9 source and amount, RNA:Cas9 ratio, repair template, and methods to enrich for edits. IDT Cas9 at 250 ng/µl was most efficient, similar to previous reports (**Figure S1A**).^1^ While RNA:Cas9 ratios from 1:1 to 10:1 have been used,^33–35^ we found that a 1:1 ratio was most efficient with our reagents (**Figure S1B**). We did not find significant improvements from 5’ modifications,^36^ (**Figure S1C**) however ssDNA templates^37^ slightly increased efficiency (**Figure S1D**). We next tested methods to enrich for edits. In pilot experiments generating collagen knock-ins we were unsuccessful with pRF4 (*rol-6(su1006)*) or *dpy-10* co-conversion^1,38^, possibly due to genetic interactions between dominant collagen mutations and the target genes. A co-CRISPR approach using a *T2A::GFP* knock-in into the 3’ end of *myo-2* yielded limited enrichment (**Figure S1E**). In contrast a pan-muscle *mlc-1p::RFP* co-injection marker enriched for edits (**Figure S1F,G**). Going forward we used this approach with unmodified dsDNA repair templates and our optimal Cas9 amount and RNA:protein ratio.

We developed a modular tagging strategy in which an initial insertion allows rapid validation of phenotype, expression, and localization. We designed an mNG::3xFLAG cassette flanked by flexible 13 amino acid linkers that contain sgRNA target sites oriented for optimal editing efficiency (**Figure 1A**).^39^ Our initial insertion includes both linker sequences and re-editing of the knock-in using the internal sgRNA sites allows for removal of the mNG::3xFLAG insertion. This design allows insertion of a desired cassette into any strain generated for the aECM toolkit using a single, common repair template and two crRNAs (**Figure 1A**). This approach ensures that our toolkit strains can be modified by any researcher in an efficient, cost-effective manner. Use of different internal and external crRNA pairs allows for removal or retention of the linker sequences during re-editing. We first tested this system by attempting to replace the mNG::3xFLAG tag in *glh-1* with an mSc::2xOLLAS tag using unmodified or biotinylated dsDNA or ssDNA repair templates. While biotinylated dsDNA and ssDNA repair templates did not boost *glh-1* knock-in efficiency (**Figure S1C,D**), they dramatically improved color swap efficiency (**Figure 1B**) suggesting variation between targets that warrants further exploration. We also validated that the other crRNA pairs allowed efficient tag swapping, except for crRNA 1+3 (**Figure 1C**). This data may suggest that crRNA #3 is a suboptimal guide and edits using it might benefit from biotinylated dsDNA or ssDNA repair templates. We successfully swapped mNG::FLAG to 2xOLLAS in 10 strains. In color swaps we also frequently observed mNG-negative animals, suggesting that the cassette was lost or inactivated (**Figure 1D**). Further optimization such as inactivation of non-homologous end-joining^40^ may be desirable. We were also able to color-swap three of the non-modular mNG alleles (*dpy-5, dpy-10, dpy-13*). As dim or stage-restricted knock-ins were more challenging to identify, we developed a complementary selection-based approach. We embedded an *unc-119(+)* selection marker in a synthetic intron in our modular mNG::3xFLAG cassette (**Figures S2, S3**) similar to a previous study,^41^ generating three strains (Figure S2B).

**Figure 1.**
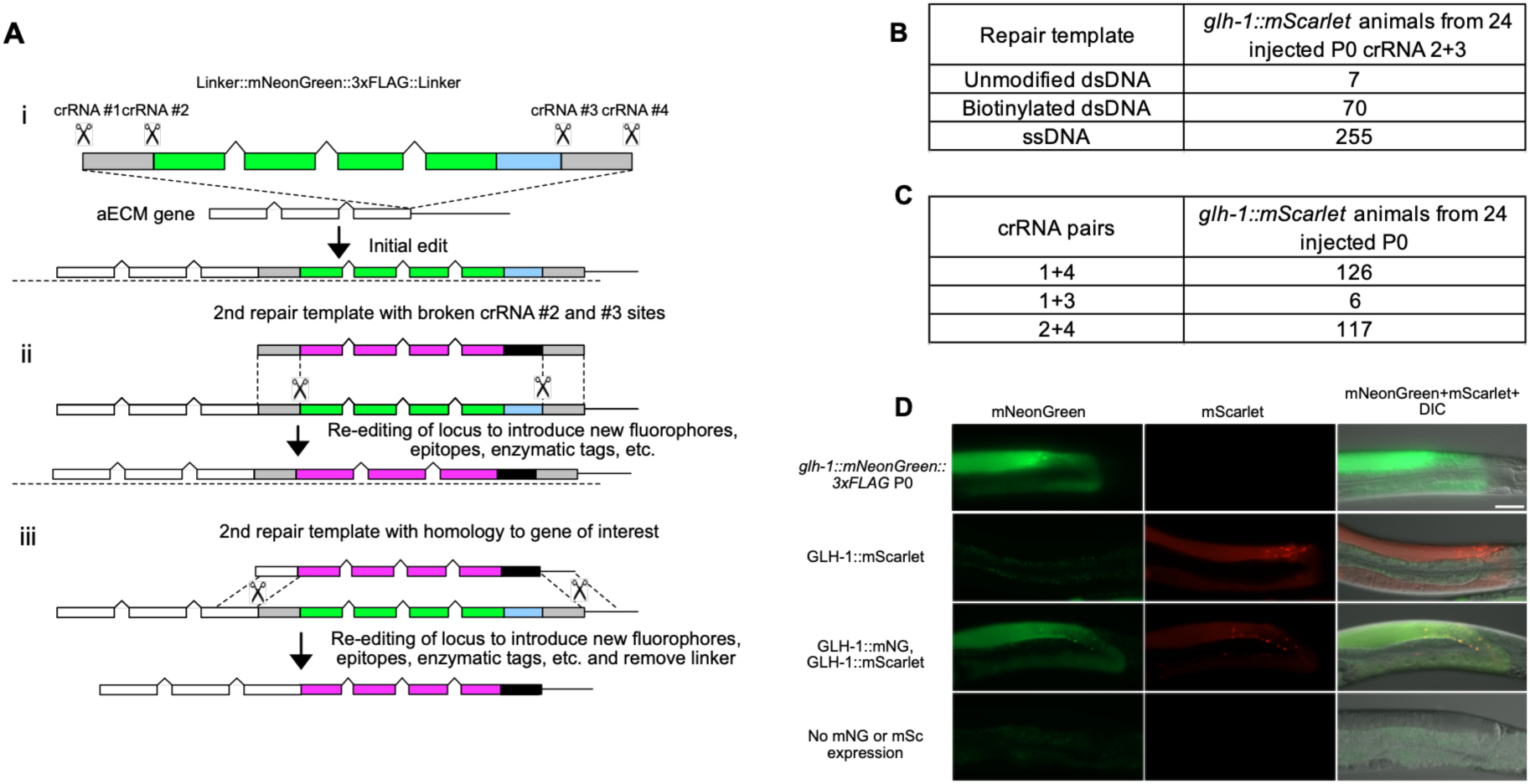
Development of a modular knock-in cassette. (A) (i) Initial knock-ins were made using an mNeonGreen::3xFLAG cassette flanked by linkers containing unique, high efficiency crRNA sites. (ii) Re-editing the initial knock-in with the internal crRNA sites (#2 and #3) allows replacement of the mNeonGreen::3xFLAG with any desired epitope. The linkers act as 35 bp homology arms and thus a single PCR repair template can be used to re-edit any initial knock-in. (iii) Using the external crRNA sites (#1 and #4) allows removal of the linker sequence during re-editing. A combination of crRNAs allows retention of the N- or C-terminal linkers during re-editing. Number of *glh-1::mScarlet::2xOLLAS* animals recovered from 24 *glh-1::mNG::3xFLAG* P_0_ animals from two independent experiments injected with the indicated repair templates and crRNAs targeting the internal sites (B) and dsDNA repair templates using the indicated crRNA pairs (C). (D) Representative images of parental *glh-1::mNG::3xFLAG* animals, and color swapped *glh-1::mScarlet::2xOLLAS* animals with and without GLH-1::mNG::3xFLAG expression. Images are representative of 20 animals imaged in two independent experiments. Scale bars = 50 µm.

### Gene families of the aECM and selection for toolkit

We focused on secreted protein families likely to comprise abundant and stable components of the aECM.^17,42^ The body cuticle is composed of collagens and non-collagenous proteins. The *C. elegans* genome contains 180 collagen-encoding genes, including the 2 divergent genes *col-55* and *col-70* (see Methods for criteria). We first focused on those for which phenotypic information existed (genetically defined collagens; bold face genes in **Table 1**). We next prioritized collagens based on expression as described above. These included collagens expressed in multiple larval stages (‘oscillating’) as well as stage-specific collagens, especially those upregulated in L4 or adult. These classes overlapped as some collagen genes are both oscillating and display stage specificity.

**Table 1.**
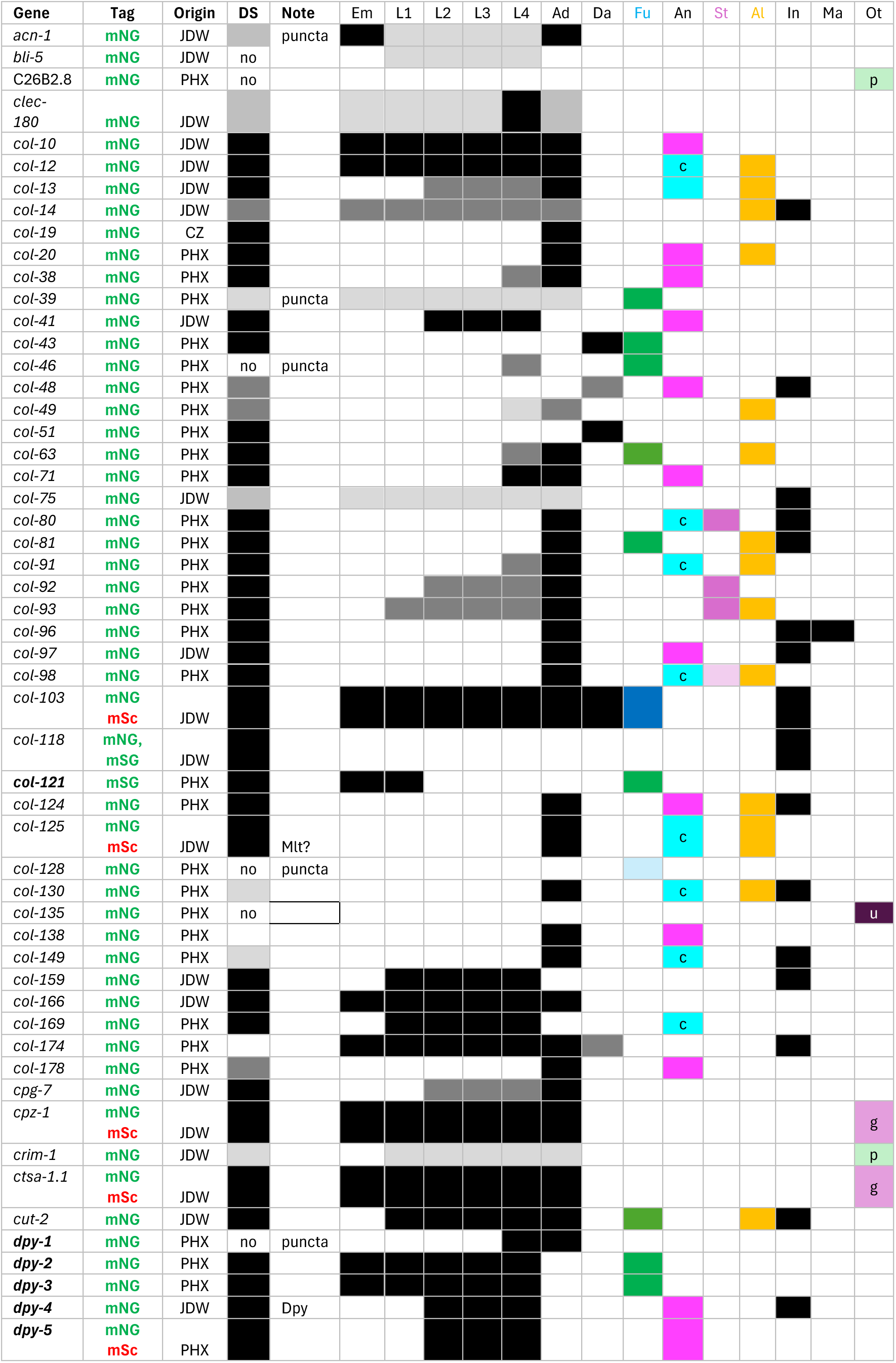
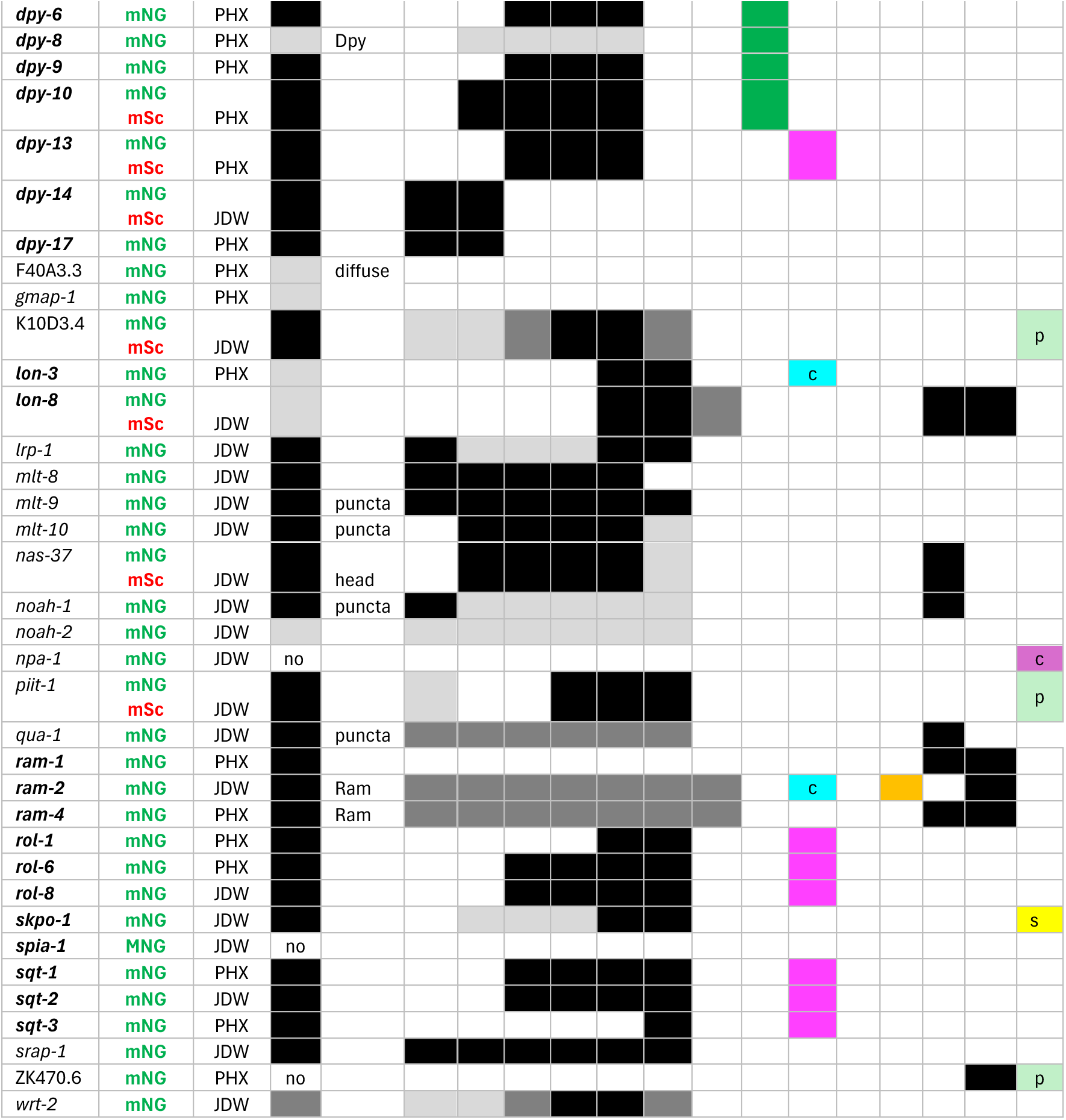
Graphical overview of aECM knock-in localization. Genes are listed sorted by gene or transcription unit name; genes first defined by mutant phenotype are in bold face. Tag: mNeonGreen (mNG) or mScarlet (mSc). Origin: Ward lab (JDW), Chisholm lab (CZ) or SunyBiotech (PHX). DS, visibility under standard dissection microscope conditions. Black box = strong, gray = weak, white = not visible. Notes: overt phenotypes or localization to epidermal puncta consistent with retention in secretory pathway. Expression and localization columns and coding: Stages: <u>Em</u>bryo, L1, L2, l3, L4, <u>Ad</u>ult, <u>Da</u>uer. aECM substructures: <u>Fu</u>rrows (green) or furrow-flanking (dark green); <u>An</u>nuli cortical (cyan, c) or basal/fibrous (magenta); <u>St</u>ruts (purple); <u>Al</u>ae (gold), <u>In</u>terfacial (sensilla, excretory pore, vulva, rectal, etc); <u>Ma</u>le (male tail rays, fan, or hook); <u>Ot</u>her: Gut (g), Pharynx (p), Uterus (u), coelomocyte (c).

While our imaging largely focused on collagens, we also tagged selected members of major non-collagen secreted protein families likely to be components of the aECM of interest to the community. We tagged the ZP domain protein DPY-1, the cysteine cradle domain proteins DPY-6^43^ and SPIA-1, ^44^ and the novel secreted protein LON-8 implicated in body cuticle morphology.^45^ We also tagged the ZP domain proteins NOAH-1 and NOAH-2, components of the pre-cuticle.^46^ We tagged the non-ZP cuticlin CUT-2 as a representative of the BME-insoluble cuticle fraction.^47^ As aECM remodeling is important in molting, we tagged astacin metalloproteases (NAS-8, NAS-37),^48^ cathepsins (CPZ-1, CTSA-1.1), and protease inhibitors (BLI-5, PIIT-1, F30H5.3, K10D3.4). We tagged CLEC-180 as a representative of C-type lectins^49^ and CPG-7 as a representative of chondroitin proteoglycans.^50^ We also tagged the mucin SRAP-1, and proteins implicated in molting (ACN-1, QUA-1, MLT-8, MLT-9, MLT-10, WRT-2).^51^

We also tagged several genes that may regulate extracellular lipids, including secreted lipid binding proteins such GM2AP/GMAP-1,^27^ the nematode polyprotein allergen related NPA-1,^52^ and the low-density lipoprotein receptor LRP-1.^53^ GMAP-1::mNG was expressed in epidermis and localized to the cuticle, consistent with prior transgenic studies.^27^ In contrast NPA-1::mNG was observed in the pseudocoelomic space and coelomocytes indicating it may be primarily secreted from other tissues. As polyprotein allergens consist of repeats that are typically cleaved into multiple small proteins, it is unclear if C-terminally tagged NPA-1 is representative of the localization of NPA-1 fragments.

### Localization of the genetically defined cuticle components

To test whether aECM components tolerate mNG tagging we first examined a set of 27 cuticle genes initially defined by mutant phenotypes.^17^ Knock-ins for four of these had been previously reported (e.g. *bli-1, bli-2, bli-6*, *dpy-13*);^7^ we generated knock-ins for the remaining genes. Knock-in strains for such genetically defined cuticle genes were morphologically normal and displayed bright fluorescence in the cuticle. The six furrow DPY collagens localized close to the morphologically defined circumferential furrows, similar to previous reports on DPY-2 and DPY-7 localization (**Figure 2A**).^44^ ^7,30,54^ Within this set individual patterns differed in detail. DPY-2::mNG showed the most consistent labeling, extending around the circumference to the lateral midline or lateral alae, without breaks (**Figure 2A**). DPY-3::mNG furrows often did not extend to the lateral midline whereas DPY-9::mNG extended to the midline but had occasional gaps (**Figure 2A**). DPY-10::mNG extended to the midline with few gaps but frequent bright dots at the ends (**Figure 2A**). Animals homozygous for DPY-2::mNG and DPY-9::mNG displayed brighter expression than single knock-in homozygotes, but with frequent gaps or dots (**Figure 2B**); neither mNG signal extended to the midline, although morphological furrows extended to the midline as visualized by DIC (**Figure 2B**). Defects in the furrow collagens affect the aECM permeability barrier function.^55^ Interestingly, BIS-1/COL-121, identified on the basis of permeability barrier defects,^56^ also localized to furrows specifically in embryonic and L1 stages. Overall, these results suggest ‘furrow’ collagen localization does not precisely correlate with morphological furrows and instead may define distinct subregions within furrows.

**Figure 2.**
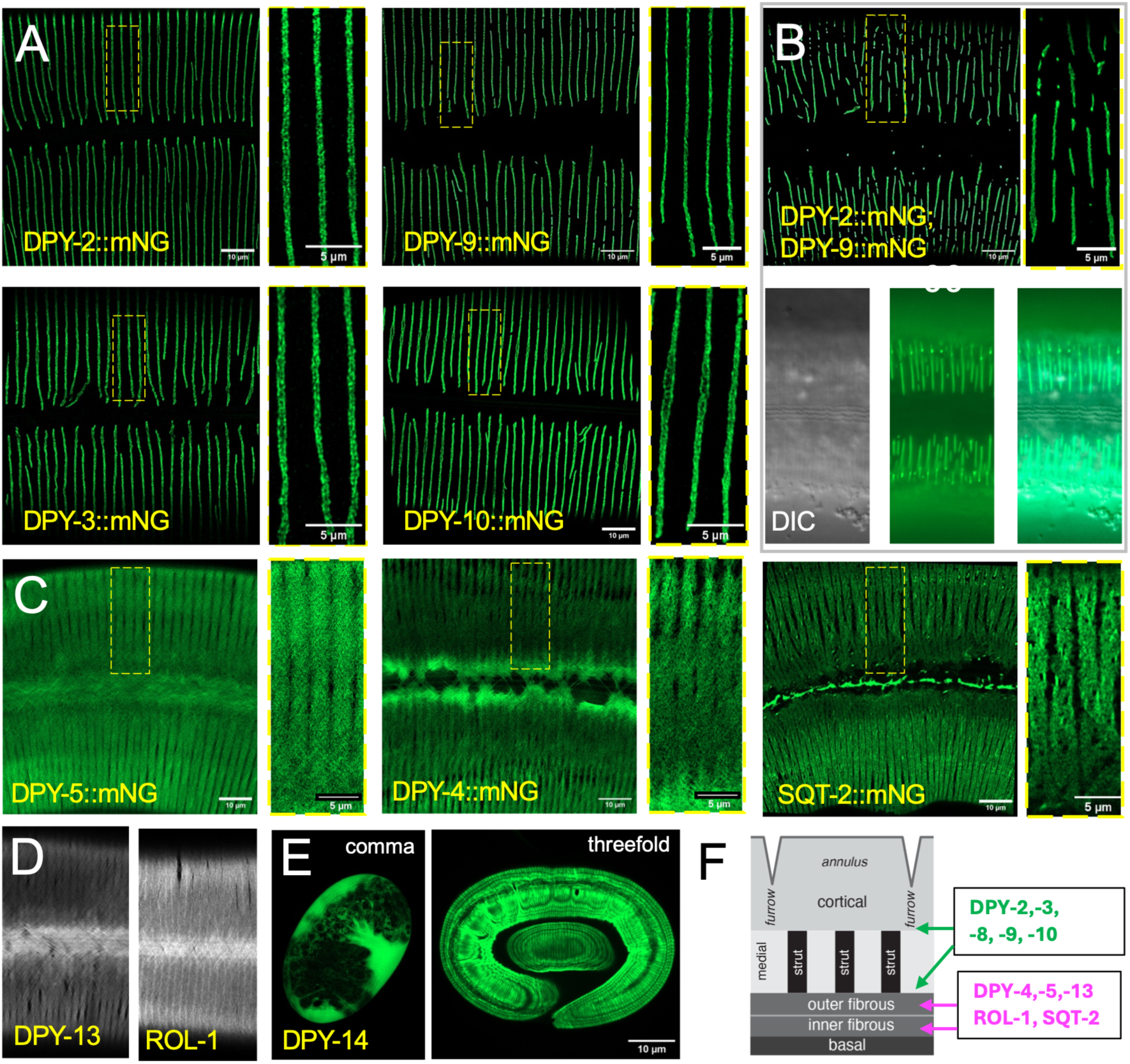
Localization of genetically defined collagens. Images are confocal z-stacks or single focal planes, oriented with anterior to the left and dorsal up; all images are of adults 24 h post L4 stage unless stated. (A) Localization of genetically defined furrow collagens DPY-2::mNG, DPY-3::mNG, DPY-9::mNG, and DPY-10::mNG. Airyscan confocal stacks. Large panels are 50 x 50 µm ROIs; enlarged insets are 12 x 6 µm. (B) Double label strain containing DPY-2::mNG and DPY-9::mNG showing fragmentation of the furrow mNG patterns. Lower panels showing DIC and widefield fluorescence. Scales, 10 µm (large panels) and 5 µm (small panels). (C) Representative images of fibrous annular collagens (DPY-4::mNG, DPY-5::mNG, DPY-13::mNG) and non-fibrous annular collagen (SQT-2::mNG). Airyscan confocal, settings as Figure 2A. (D) DPY-13::mNG and ROL-1::mNG localization in adult. (E) DPY-14::mNG in comma and threefold stage embryos. Scales, 10 µm (large panels) and 5 µm (small panels). (F) Cartoon cuticle cross section indicating furrow and fibrous/basal collagen localization for collagens expressed in adult cuticle. There is evidence for furrow collagens in the basal layer, but it is unclear whether they also localize to the cortical layer. Longitudinal aspect showing 1 annulus, two furrows, and three struts.

Two other collagen knock-ins, DPY-5::mNG and DPY-13::mNG, were enriched in annuli (**Figure 2C,D**), forming crossed fibrous arrays corresponding to the two fibrous layers of the cuticle;^57^ we classified these and similar patterns as ‘annular fibrous’. DPY-4::mNG displayed a more braided fibrous pattern within annuli (**Figure 2C**). These three collagens were not expressed in embryos but were upregulated in late L1 and visible as fibrous patterns in L2 onwards (**Figure S4**), consistent with the lack of fibrous layers in L1 cuticle.^58^ DPY-14::mNG and DPY-17::mNG were expressed beginning in embryos and localized to annuli in the L1 cuticle (**Figure 2E**), consistent with the early larval Dpy phenotypes of *dpy-14* and *dpy-17* mutants. LON-3::mNG localized to double bands within annuli but did not show overtly fibrous organization. Three collagens originally defined by male <u>r</u>ay <u>a</u>bnormal <u>m</u>orphology (Ram) phenotypes^59^ displayed distinct localizations. RAM-1/COL-64::mNG was only detected in male rays and the male hook cuticle. RAM-2/COL-82::mNG localized to annuli and lateral alae (longitudinal ridges) of both sexes as well as the male tail fan. RAM-4/COL-34::mNG localized to interfacial cuticles in both sexes and the male tail fan. *lon-8*, which also displays Ram phenotypes,^45^ encodes a novel secreted protein; LON-8::mNG localized to the cuticle of the vulva (**Figure S5**) and male tail fan, as well as to adult and dauer alae. These observations suggest the Ram phenotype can result from aberrant function of multiple proteins in the male tail aECM. Knock-ins for ROL-1 and SQT-2 collagens localized to annuli (**Figure 2B,C**), among which ROL-1::mNG and ROL-6::mNG were the most fibrous and SQT-1::mNG the least. Rol phenotypes may be due to direct or indirect disruption of the fibrous layers.^60^ Many of the above knock-ins were readily detectable in shed cuticles, consistent with their accumulation in aECMs shed at molts. Overall, these findings suggest the genetically defined collagens are primarily localized to furrows or to the fibrous layers (**Figure 2F**).

In the course of this analysis we found that *sc22,* annotated as a *rol-1* allele,^60^ did not contain a lesion in *rol-1* and was weakly linked to the ROL-1::mNG knock-in. Analysis of the *sc22* strain sequence revealed two other SNPs on chromosome II that would result in collagen missense alterations, including one causing R106H (RTAR > RTAH) in the *col-71* dibasic cleavage site. A *col-71(R106H)* mutation edited into a wild-type background also caused an adult Rol phenotype, suggesting this mutation is causative. Notably, the *rol-1* reference allele *e91* also has a dibasic cleavage site alteration (RTAR > RTAC).^61^ Like ROL-1::mNG, COL-71::mNG displayed an annular fibrous localization (**Figure 4A**), consistent with it playing a similar role to ROL-1 in the adult cuticle.

An important question at the outset was whether collagens readily tolerate tagging. *C. elegans* cuticle collagens undergo N-terminal cleavage by subtilisin proteases and otherwise largely consist of Gly-X-Y repeats that are critical for function. Thus, we tagged most collagens at their C-termini for convenience; in a small number of cases we inserted a tag within the protein. For example, SQT-3 is cleaved at the C-terminus ^62^ therefore we inserted a tag internally ^63^. Several other collagens we tagged contain similar C-terminal cleavage sites, however C-terminal knock-ins displayed cuticle localization. Of the cuticle collagens defined by mutant phenotype we detected phenotypes for only three knock-in strains. The DPY-4::mNG and DPY-8::mNG KI strains displayed mild Dpy phenotypes whereas RAM-4/COL-43::mNG and RAM-2/COL-82::mNG KI males displayed Ram phenotypes, suggesting these insertions affect gene function. The SQT-1::mNG knock-in allele suppressed the Bli phenotype of *bli-2(0)*, consistent with it impairing *sqt-1* function^7^. As mentioned above, animals homozygous for DPY-2::mNG and DPY-9::mNG knock-ins displayed fragmented furrow collagen localization (**Figure 2B**) but were not overtly Dpy. In contrast, animals homozygous for two fibrous layer knock-ins DPY-5::mSc and DPY-13::mNG resembled the single homozygotes in expression and localization. Other strains combining two knock-ins did not display overt interactions suggesting synthetic effects may be specific to certain knock-in combinations. Although this analysis was not exhaustive it suggests most cuticle collagens tolerate C-terminal knock-ins; in some cases tagging may result in cryptic loss- or reduction-of-function that can be detected in sensitized backgrounds.

Most of our analysis relied on homozygous knock-in strains. To assess how tagged collagens compete with untagged proteins in the heterozygous state we examined a set of 7 collagen knock-ins expressed at varying levels. Interestingly, the fluorescence intensities of cuticle mNG of most knock-in heterozygotes (5/7) were significantly above 50% that of the homozygotes; 2/7 (*col-81*, *sqt-2*) were not significantly different from 50% of homozygote intensity (**Figure S5A**). With the caveat that we have not assessed protein levels, these comparisons suggest tagged collagens can be incorporated as efficiently into the ECM as untagged alleles. Most KIs exhibited minimal recovery in FRAP experiments except for LON-8::mNG in the L4 vulva (**Figure S5B**), consistent with tagged proteins being stably incorporated into the matrix.

Many knock-ins were visible within underlying epidermal cells, localized to puncta that likely correspond to secretory organelles^7^ (’punctate’ in **Table 1**). Some mNG-tagged collagens accumulated in puncta within the epidermis and showed little cuticle fluorescence; in these cases, tagging may have interfered with secretion and the mNG-tagged proteins may be aberrantly retained in secretory compartments. Some mSc-tagged knock-ins showed intense fluorescence in filamentous structures that resemble epidermal lysosomes.^64^ It is unclear if these reflect normal endocytosis or aberrant trafficking of tagged proteins.

### Other collagens and cortical aECM compartments

In addition to the above genetically defined collagens, we tagged an additional ∼31 cuticle collagens and a non-cuticle collagen (*col-135*). As mutant phenotypes were not known for most of these, functionality of the tagged proteins could not be verified. All were detectable using dissecting or widefield microscopy. While some showed patterns reminiscent of genetically defined collagens, many showed distinctive localization especially to cortical layers. We summarize these patterns by compartment:

#### Furrow or Furrow-Flanking

Four other collagens consistently localized to furrow-like patterns (**Figure 3; Figure S6**) resembling the six known ‘furrow’ collagens, although none were as specific as the furrow collagens. These new furrow-localizing or furrow-flanking collagens included COL-46::mNG (also substantially in epidermis), COL-103::mNG (thick furrows), and COL-149::mNG (punctate along furrows and in annuli). Two proteins showed distinctive ‘furrow-flanking’ localization to double bands immediately adjacent to furrows defined by DIC: the cuticlin CUT-2::mNG and the adult specific collagen COL-63::mNG (**Figure 3B,C**). The COL-63::mNG double-band pattern was reminiscent of the localization of BLI-1::mNG and BLI-2::mNG knock-ins to furrow-flanking bands prior to their condensation into punctate struts,^7^ however COL-63::mNG double bands persisted into adulthood, suggesting they define a distinct compartment of the adult furrow substructure. In *dpy-3(e182)* mutants COL-63::mNG and CUT-2::mNG had a punctate or vermiform appearance (**Figure 3B**), indicating their localization is dependent on normal furrow formation.

**Figure 3.**
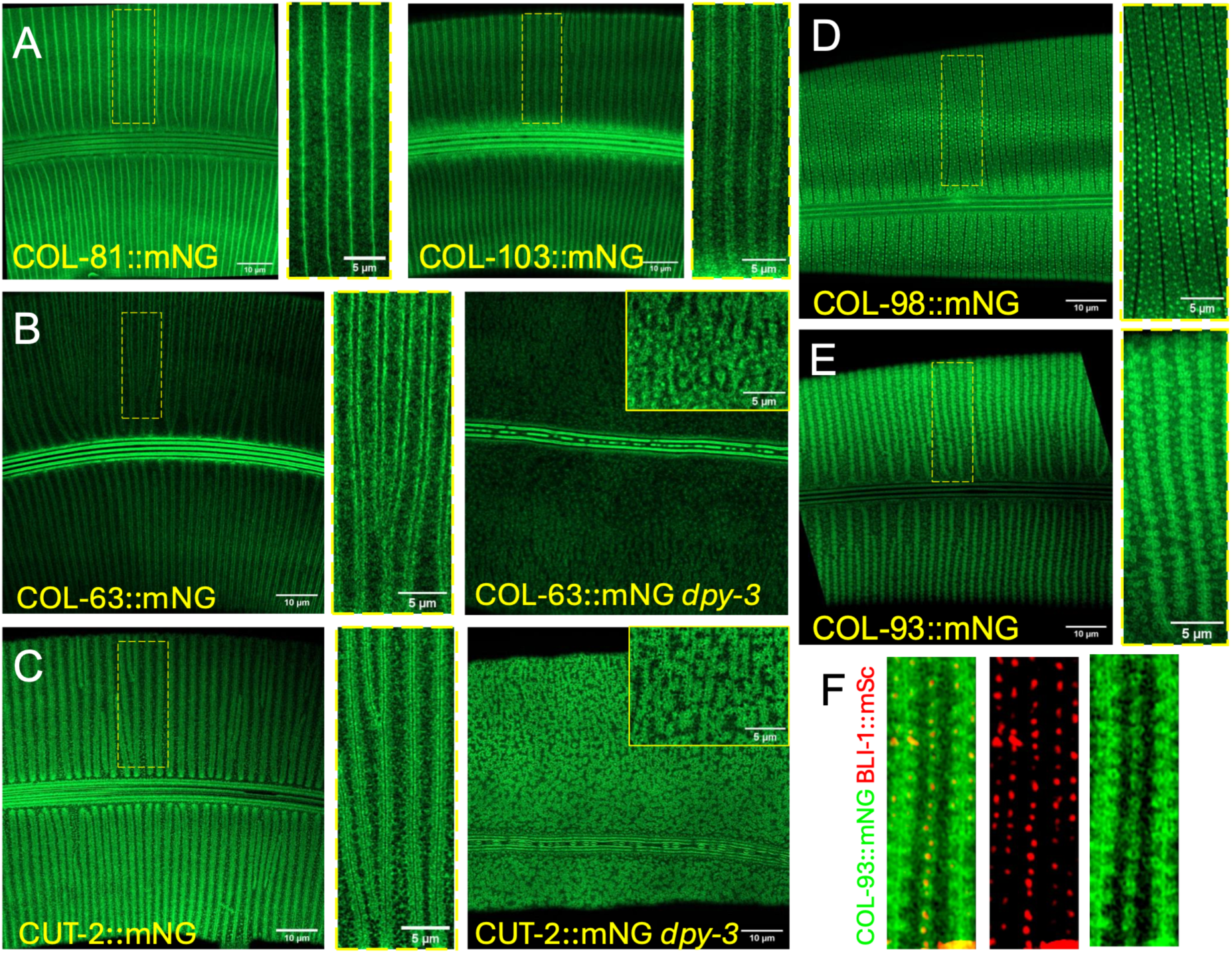
Novel Furrow, Furrow-flanking, and Strut-related patterns. (A) Airyscan confocal projections of COL-81::mNG and COL-103::mNG showing localization to furrow or furrow-flanking patterns, based on spacing and orientation relative to morphological furrows. (B) COL-63::mNG localization to furrow-flanking pattern. In *dpy-3* mutants the COL-63::mNG furrowflanking pattern was converted to a dispersed punctate pattern. (C) CUT-2::mNG localization was similarly aberrant in *dpy-3* mutants but was less punctate than COL-63::mNG. Airyscan confocal. Scales, 10 μm (large panels) and 5 μm (small panels). (D, E) Airyscan confocal projections of COL-98::mNG and COL-93::mNG (Airyscan). COL-103::mNG also localizes preferentially to furrow-flanking struts, more evident in basal sections (See **Figure S3**). (F) Colocalization of COL-93::mNG and BLI-1::mSc. COL-93::mNG circles surround BLI-1::mSc puncta in a z-projection (left panel). Scales, 10 or 5 μm. Panels in F are 4 μm wide.

#### Struts

Struts are column-like connections between basal and cortical layers, formed in late L4 and containing three collagens (BLI-1, BLI-2, BLI-6)^7^ and two epicuticlins EPIC-1 and EPIC-2.^21^ We found several additional collagens that localized to adult-specific puncta corresponding to struts (**Figure 3; Figure S7**), although none were exclusive to struts. COL-98::mNG displayed faint strut-like puncta as well as diffuse localization in annuli (**Figure 3D; Figure S7**). COL-93::mNG showed a unique localization to rings around strut bases (**Figure 3E,F; Figure S7**). Compared to BLI-1 or EPIC-1-labeled strut ‘donuts’, the COL-93::mNG rings were larger (0.44 ± 0.07 µm diameter, mean ± SD, n = 122) and slightly more basal than BLI-1::mNG (**Figure 3F**). COL-93::mNG and COL-98::mNG were diffuse in the late L4 cuticle, so like the EPIC proteins may be recruited to struts after the L4-adult molt. Similarly, COL-80::mNG localized to furrows in the L4 stage; the strut-like puncta developed gradually over first 48 h of adulthood. COL-103::mNG and COL-81::mNG showed weak strut localization in addition to furrow localization (**Figure 3A, Figure S5**) suggesting struts and furrows may have overlapping composition. Our observations suggest struts are intricately patterned substructures with at least 9 distinct protein components.

#### Fibrous patterns

Several collagens displayed ‘annular fibrous’ patterns resembling those of the genetically defined collagens DPY-4/5/13 or ROL-1 (**Figures 2D, 4A**). The adult specific collagen COL-19 was one of the first to be GFP-tagged in transgenics^65^ but details of its localization had not previously been analyzed. COL-19::mNG localized to oriented fibrous layers^7^ and to less consistently oriented fibers deep in the lateral cuticle. COL-38::mNG and COL-138::mNG both displayed adult-specific annular fibrous organization. Notably, transcription of both *col-38* and *col-138* is activated in the late L4 stage by the zinc finger transcription factor LIN-29, an important heterochronic pathway regulator.^66,67^ Although the function of these adult-specific fibrous collagens is not known, they may contribute to adult cuticle strength.

Most such ‘annular fibrous’ collagens localized to the crossed fiber arrays. By examining COL-71::mNG in closely spaced focal planes (**Figure S8**; see Methods) we determined that the outer (more cortical) fibers formed right-handed helices whereas the inner array formed left-handed helices, consistent with earlier EM observations.^57,68^ In the midbody, the angle of these fibers in either array to the long axis varied along the dorsoventral axis, with lateral fibers forming shallower angles (30-40°) compared to more dorsal or ventral regions (up to 80°) (**Figure 4B**), whereas in the anterior head the arrays were more consistently angled at ∼65° to the long axis. Although most fibrous collagens were in both the outer and inner fibrous arrays, some appeared to selectively label a single array. For example COL-20::mNG appeared predominantly in the outer right-handed array as well as to curving oriented fibers deeper in the cuticle (**Figure 4A; Figure S8**); the latter may represent the basal-most fibrillar layer. Some collagens displayed fibrous patterns distinct from the crossed fiber arrays (**Figure 4C**). These included COL-48::mNG, which localized to circumferentially oriented bundles of fibers, and COL-178::mNG, which localized to long thick curved fibers (**Figure 4C)**. COL-41::mNG localized to annular patterns in L2-4 as well as to oriented fibers in the lateral cuticle.

**Figure 4.**
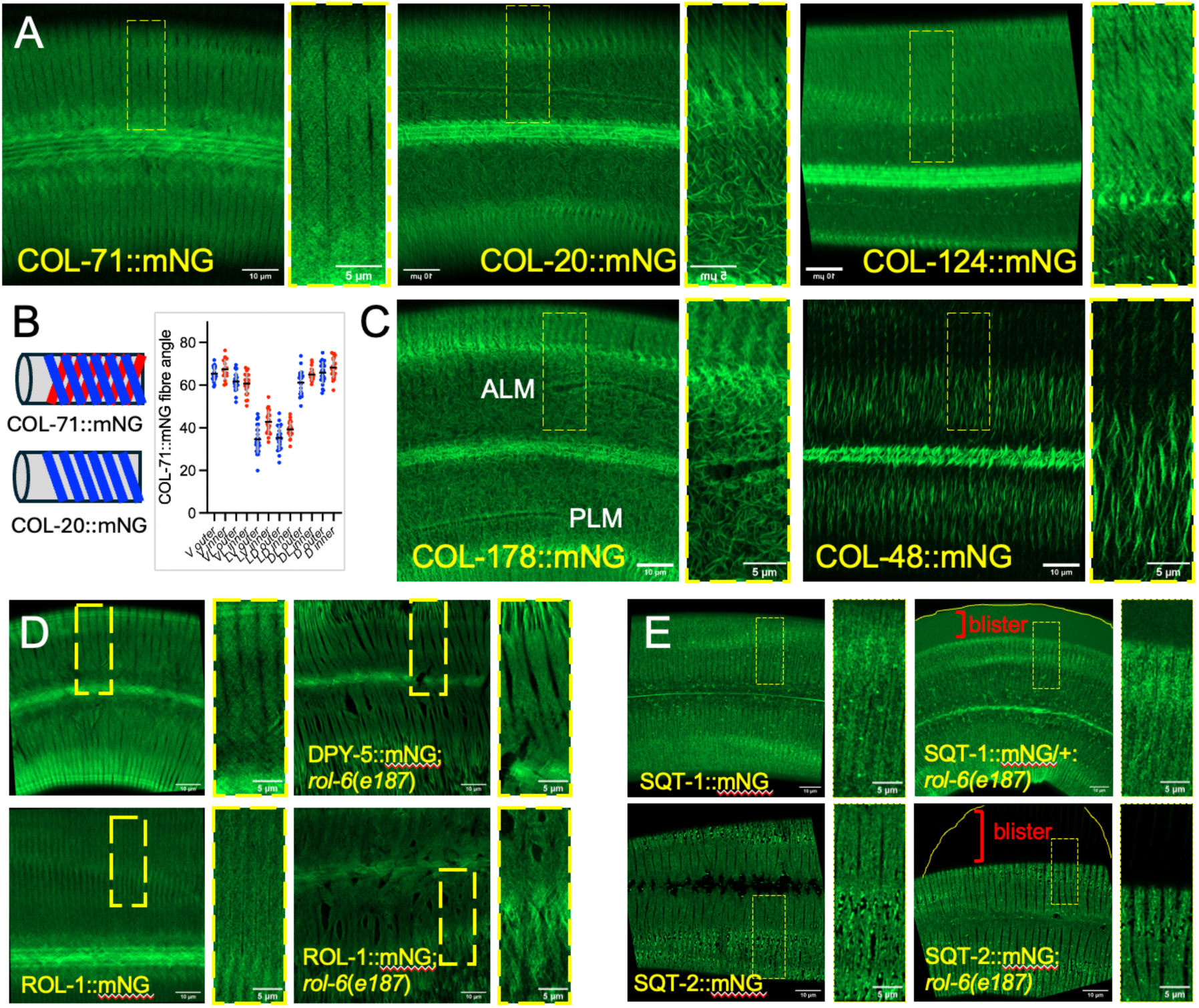
Markers for the oriented fibrous arrays in basal layers. (A) Airyscan confocal projections showing fibrous layer localization of COL-71::mNG (both layers; single confocal slice), COL-20::mNG (inner layer), and COL-124::mNG. Deeper COL-20::mNG fibers (more lateral epidermis; lower part of inset) are not consistently oriented. (B) Cartoon of fibrous array chirality. The outer fibrous layer forms as a right-handed helix (blue); the inner layer is left-handed (red). COL-71::mNG is found in both layers whereas COL-20 is predominantly in the outer layer based on its right-handed helical pattern. Quantitation of angle of fibrous arrays to long axis at different points in dorsoventral axis in outer and inner layers, measured for COL-71::mNG in midbody. Mean ± SD, n = 20-25 for each set. (C) Other examples of fibrous arrays: COL-178 forms dense filigreed patterns in the basal cuticle, as well as faintly localizing to oriented fibrous layers. COL-48::mNG forms short circumferentially oriented fibers, especially under lateral alae. (D) Fibrous layer markers DPY-5::mNG and ROL-1::mNG are mislocalized in *rol-6(gf)* animals. (E) SQT-1::mNG or SQT-2::mNG markers cause Bli phenotypes in *rol-6(gf)* backgrounds. Scales, 10 µm or 5 µm (insets).

Mutations in multiple collagens result in ‘Roller’ (Rol) phenotypes in which the cuticle and underlying tissues are helically twisted, potentially due to aberrant fibrous layers. ^60^ We therefore examined fibrous layer markers in Rol mutant backgrounds. Such markers were variably mislocalized in *rol-6(gf)* backgrounds (**Figure 4D,E**), however crossed fibers remained visible (**Figure 4D**), suggesting Rol mutations may indirectly affect fibrous layers. Interestingly, introduction of SQT-1 or SQT-2::mNG in Rol background frequently resulted in Bli phenotypes (**Figure 4E**), suggesting these KIs have cryptic effects on function. Speculatively, such combinations could impair fibrous layer attachment to struts.

#### Annular and diffuse localization

Several collagens showed annular localization but were not overtly fibrous (**Figure 5**). COL-91::mNG displayed a slightly granular localization within annuli, reminiscent of EPIC-1 or EPIC-2 proteins; COL-125::mNG and COL-130::mNG displayed a smooth annular localization. COL-81::mNG localized to annuli in L4s and males but was more furrow-like in adult hermaphrodites. Some knock-in strains displayed localization that was less easily classified due to their low fluorescence level (e.g. COL-39::mNG, COL-49::mNG) but may be diffusely localized within multiple compartments.

**Figure 5.**
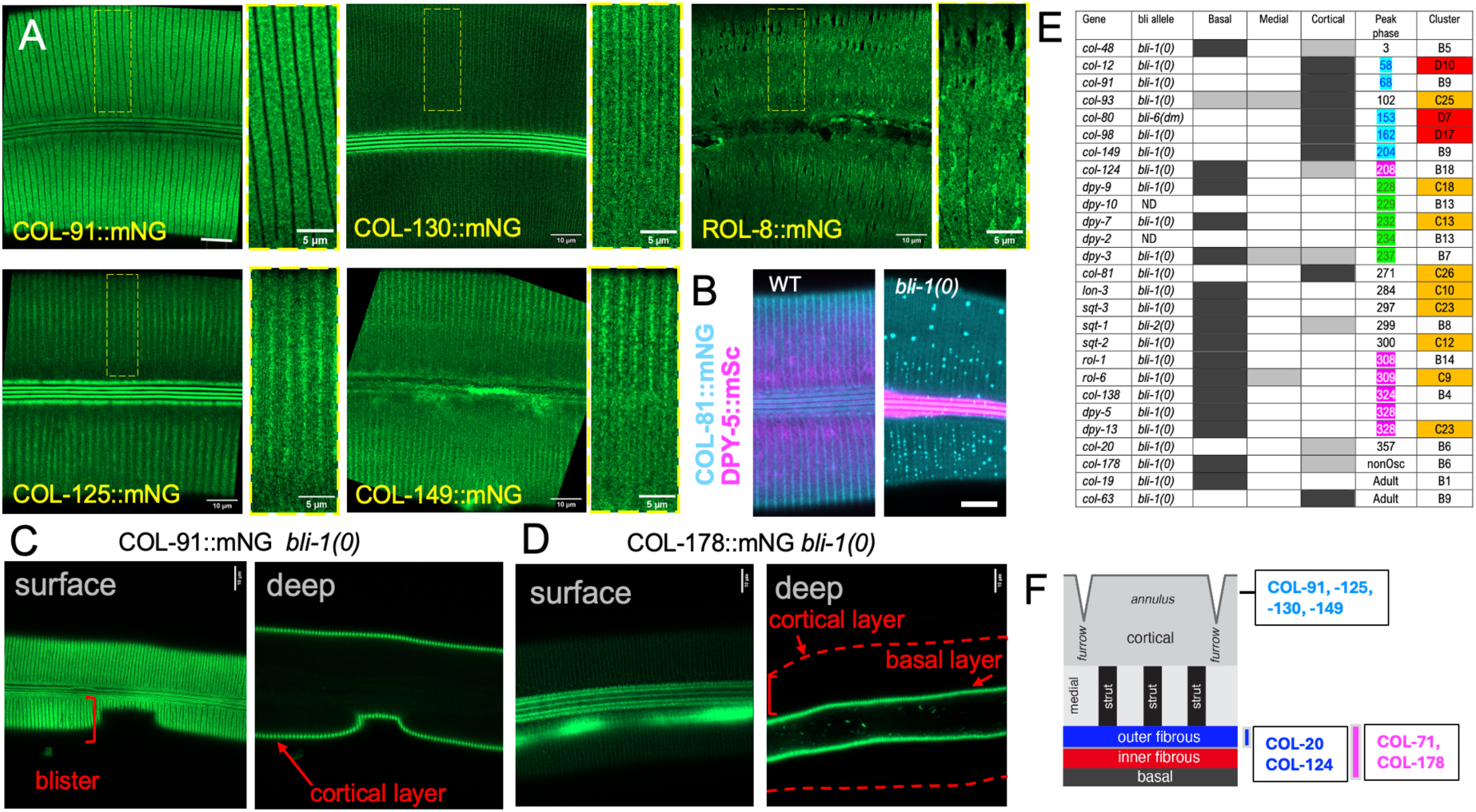
Non-fibrous annular and cortical compartments. (A) Airyscan confocal projections showing annular localization patterns: COL-91::mNG is diffusely localized throughout annuli. COL-125::mNG is concentrated in the central region of annuli. (B-D) Cortical versus basal localization in blistered mutants. B, Double labeling of COL-81::mNG and DPY-5::mSc in wild type and *bli-1(0)* mutants. In *bli-1* mutants COL-81::mNG remains on the cortical surface and becomes punctate; DPY-5::mSc is only found basally. Scale, 10 µm. C, Surface and deep confocal sections of COL-91::mNG (C ) and COL-178::mNG (D) in *bli-1(ju1395)* null mutants. Blistered regions indicated with square brackets. COL-91::mNG localizes exclusively to the outside of blistered regions (cortical) and COL-178::mNG localizes exclusively to the inside (basal). Scales, 10 µm. (E) Table of cortical versus basal localization Gray shading indicates strong, weak or no fluorescence in the respective layer. Phase angles for oscillating collagens are based on ^3^; approximate phase angles for lethargus entry and ecdysis are from ^4^; nonOsc, transcript not oscillating; Adult = adult-specific. Cyan lettering indicates cortical localization, green indicates furrow localization, and magenta indicates fibrous layer localization. We interpret ‘cortical’ collagens as the earliest wave of collagen expression for the subsequent cuticle, however as timing of protein expression to ecdysis remains unclear, they may represent a late wave.^6^ Localization of DPY-7::sfGFP and COL-19::mNG in *bli* mutants has been previously reported.^7^ Collagen clusters are based on domain organization and do not strictly correlate with sequence identity;^9^ cluster D are in red and C in orange. (F) Cartoon summary of cortical and fibrous layer localizations from Figures 4 and 5.

#### Cortical versus basal compartments

To assess whether a specific knock-in fusion protein localized to cortical or basal compartments we used Bli mutants in which cortical and basal layers become widely separated in adults. We previously used this approach to show that COL-19::mNG and DPY-7::mNG both localized basally in blistered animals.^7^ We examined an additional 24 knock-ins in Bli mutant backgrounds including 23 collagens and DPY-6. Using this assay many collagens could be classified as exclusively cortical (7) or exclusively basal (9); a minority localized to more than one layer in *bli* mutants (**Figure 5 E**). We generated further double-label strains using cortical mNG-tagged knock-ins and the basal marker DPY-5::mSc, and found that markers such as COL-81 consistently localized in more cortical focal planes to DPY-5::mSc in *bli-1(0)* (**Figure 5B**). Cortical collagens such as COL-91:mNG retained their annular patterning in *bli-1(0)* (**Figure 5A,C**); basal collagens such as COL-178::mNG remained in annular fibrous patterns in *bli-1(0)* (**Figure 5D**). In contrast, COL-80::mNG strut localization was abolished in *bli-1(0)* mutants and the remaining COL-80:mNG variably localized to annuli or furrows. Thus, in some cases, disruption of struts may affect cuticle organization beyond separation of cortical and basal layers.

Cuticle collagen transcript levels oscillate during larval development, with different collagens showing distinct peaks at different phases during each larval stage (see Introduction) ^3,4^. *dpy-6* is one of the first genes expressed in the molting cycle (peak phase 165°) whereas *nlp-29* is one of the last.^44^ Cortically localized proteins correspond to transcript peak phase angles between 50-200° (**Figure 5E**) whereas basal collagen transcripts peaked at later phases (e.g. transcripts of the 6 *sqt* or *rol* collagens peak between 290-310°). This timing would suggest the cortical collagens are expressed either just before the molt, presumably for construction of the cuticle early in the next larval stage, or right after the molt. As it is unclear how cortical collagens would be assembled outside basal layers, we favor the interpretation that the cortical collagen genes are expressed very early in the cyclic synthesis of each aECM. Interestingly, many basally localized collagens have been identified based on morphogenetic mutant phenotypes, whereas cortically localized collagens have not been found in phenotype-based screens (**Figure 5F**).

#### Lateral Alae

Several knock-ins localized to alae, the cuticular ridges that extend longitudinally along the lateral cuticles of L1s, dauer, and adults;^29,69^ examples are in **Figure S9A**. Adult alae localization could be broadly classified as localization to the three alae ridges (e.g. COL-125::mNG, COL-130::mNG), to the flanks of ridges (six longitudinal stripes; e.g. COL-103::mNG; COL-63::mNG), or to the valleys between the ridges (2 or 4 stripes; e.g. COL-81::mNG, CUT-2::mNG). Some collagens localized to deeper fibers beneath alae (e.g. COL-14::mNG). Several collagens also localized to the alae of L1 or of dauers (**Figure S9B,C**).

#### Specialized aECMs of interfacial cuticles and other tissues

In addition to the body cuticle compartments defined above, many collagens showed strong expression in specialized interfacial aECMs such as those of the nose tip (including the mouth and amphid and other sensilla), vulva, rectum, excretory pore/duct, or pore-forming sensilla such as the phasmids and deirids (**Figure 6A**; **Table 1**). For example, COL-34::mNG and COL-174::mNG were detected in many interfacial cuticles (**Figure 6B,C**). localized exclusively to male tail rays, whereas RAM-2/COL-82::mNG and RAM-3/COL-34::mNG also localized to the fan aECM and caused Ram phenotypes (**Figure 6D**). Numerous collagens localized to distinct regions of the vulval cuticle (**Figure 6E**). These observations reinforce findings that interfacial aECMs are highly specialized due to local secretion of aECM^26^ ^70^ or local determinants of assembly.^71^ Although COL-51, COL-92 or SQT-2 have been hypothesized to function in an intestinal aECM, we did not detect intestinal localization in these or other collagen KI strains. ^72,73^ Indeed, all tagged collagens were found in epidermis or cuticle, apart from the divergent collagen COL-135, which was expressed in spermatheca and uterus (**Figure S10**) and therefore may be a component of the little-studied uterine matrix.^74^

**Figure 6:**
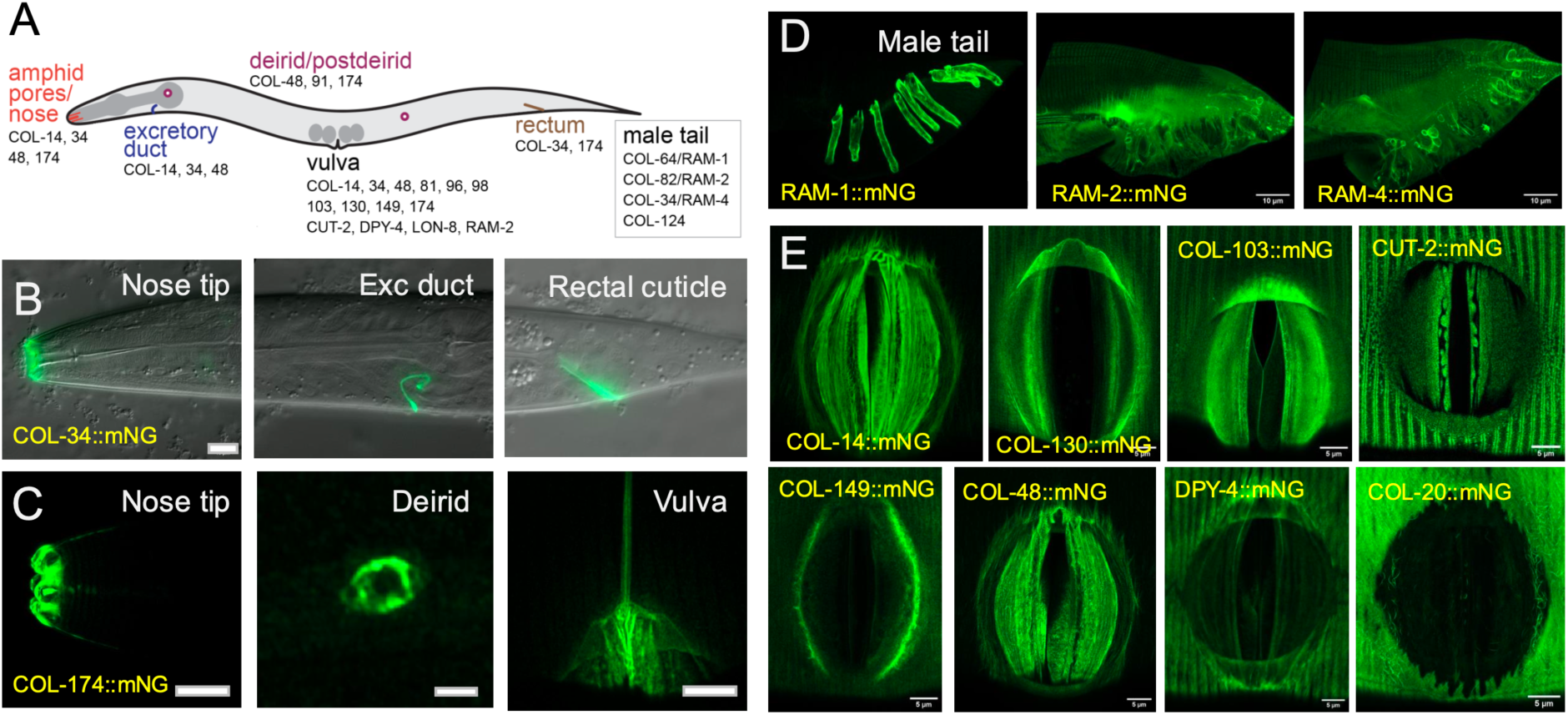
Localization of collagens and non-collagens to interfacial aECMs. (A) Cartoon of interfacial aECMs in adult hermaphrodites, showing nose tip (mouth, buccal cavity, amphid pores), excretory pore and duct, vulva, rectum, deirid and postdeirid. Localizations of selected knock-ins are indicated; see also **Table 1**. (B) COL-34:mNG (RAM-4) localization to nose tip, excretory duct and rectal cuticle, widefield fluorescence and DIC. (C) COL-174::mNG localizes specifically to interfacial cuticles; shown is localization of the nose tip, deirid and vulva (lateral view) Scales, 10 µm for nose tip and vulva, 2 µm for deirid. (D) Localization of RAM collagens (COL-64, COL-82, COL-34 to male tail ray and fan aECMs. (E) Localization of COL-14, COL-130, COL-103, CUT-2, COL-149, COL-48, DPY-4, and COL-20 mNG knock-ins to vulval cuticle in early adult, ventral views; Airyscan confocal projections. Scales, 10 µm (B,D), 5 µm (E).

## Discussion

We report the first large scale set of endogenously tagged aECM components in any organism. Our set of molecular markers reveal the exquisite tissue and stage specificity of aECMs, and the multiscale patterning of aECM substructures. Our work has implications for CRISPR methodology, for the molecular architecture of the *C. elegans* aECM, and for the spatiotemporal organization of complex ECMs.

### Molecular architecture of apical ECMs and regulation of aECM expression

Among the *C. elegans* aECMs, the cuticle alone likely contains hundreds of components. We consider whether, comparable to BM, the cuticle aECM may be dissected into ‘core components’ expressed at all or most stages and cuticle types, on top of which are added stage and region-specific elements. Of the genetically defined cuticle components, some were expressed in all stages, and may form a ‘core’ cuticle aECM to which additional stage or structure specific components are added. These include components of annuli such as SQT-3, and several furrow collagens, consistent with the presence of annuli and furrows in all cuticle types. However, our gene set was slightly biased toward collagens expressed in a subset of larval stages; of ∼20 collagens expressed at all stages by RNAseq we have only tagged SQT-3, COL-12, and RAM-4/COL-34, which is found in interfacial cuticles.

Clear examples of stage-specific components include the L1 collagens DPY-14 and DPY-17, the L2-4 stage collagen COL-41, and the numerous ‘adult specific’ collagens. It should be clarified that stage-specific collagens are first expressed in the stage preceding the stage in which they are incorporated into matrix: i.e. many ‘adult specific’ collagens are first transcribed in L4, secreted in late L4, and form stable structures in the adult cuticle.^7^ Our knock-in analysis was consistent with the L4 upregulation of four other collagens known to be targets of the LIN-29 transcription factor.^66^ Some aECM transcripts are first detected in L4 and are present at higher levels in adults, whereas others are first detected in adult. Interestingly, some genes such as *col-12* show an additional transcript peak early in adulthood^6^ and our protein analysis was consistent with this observation. The mechanistic basis of the early adult peak is not known but could be related to larval transcript oscillations. Several proteins such as EPIC-1 ^21^ display distinct localization in adult versus L4 cuticle, suggesting the adult cuticle undergoes maturation well beyond the L4/adult molt.

We found several collagens that localize exclusively to the cortical compartment of the cuticle. In contrast, most genetically defined collagens localize to more basal or fibrous layers. The cortical compartment is an exciting area for investigation as it may contribute to surface properties involved in host-pathogen interactions or mate recognition.^25^ Equally, the architecture of the fibrous layers is little-understood. Crossed fibrous arrays are integral to the strength of the hydrostatic skeleton in many nematodes and other animals,^75^ however little is known about their biogenesis. Our markers reveal that the crossed fibrous arrays are variably angled with respect to the long axis, reminiscent of variable-angle composites that maximize load-bearing capacity.^76^ Moreover, we identified collagens unequally distributed between the two fibrous layers. Aberrant patterning of such fibrous layers has long been hypothesized to underlie helically twisted ‘Rol’ mutant phenotypes.^60^ Our resource provides tools to address how fiber angle is regulated and how the two fibrous arrays are laid down with consistent chirality.^68^

The dauer cuticle is highly specialized, with unique structures such as a basal nanoscale mesh that may contribute to stress resistance. Transcriptomic studies show ∼30 cuticle collagens are transcribed predominantly in dauer or predauer stages.^77^ We did not prioritize such dauer-specific collagens as their low expression in non-dauer stages made them more challenging for fluorescence-based screening. Two dauer-specific collagens tested, COL-43 and COL-51, indeed were only visible in dauer and/or predauer cuticle. Additionally, several proteins expressed in other stages showed distinctive localization in dauer larvae, e.g. COL-48 (dauer deirids) or LON-8 (dauer alae). An important future direction would be to focus on the dauer-specific aECM and our *unc-119(+)* pipeline may be best suited for these genes.

### Collagen localization determinants and timing

Our analysis indicates that collagen localization in the aECM shows a high degree of specificity, in that almost every collagen knock-in showed a slightly different pattern. Most *C. elegans* cuticle collagens are 300-400 aa; after cleavage, mature cuticle collagens mostly consist of Gly-X-Y domains and short N- and C-terminal regions. Cuticle collagens have been classified in several ways, such as the patterns of interruptions in the Gly-X-Y domain and presence of a signal peptide or transmembrane domain.^9^ It remains unclear which sequence features might direct localization.

Interestingly, of four tagged collagens classified in cluster D ^9^, three were cortical (**Figure 5E**). Our toolkit provides a resource for the dissection of determinants of collagen localization.

Timing of aECM protein expression could provide additional information. Within each larval stage collagen transcription occurs in waves,^6^ resulting in oscillating transcript levels.^32^ Such observations led to models that the cuticle could be built in successive layers, with early-peaking proteins forming outer layers and later-peaking proteins forming inner layers. Such models are an oversimplification given the early expression of precuticle proteins^17^ and the lag between collagen secretion and aECM assembly in the embryo.^63^ Nevertheless, our analysis suggests that cortically localized proteins tend to have early transcript peaks whereas basal proteins have later peaks. Models whereby waves of collagen deposition add cuticle layers from outer to inner may account for aspects of cuticle layer formation.

For the current toolkit we prioritized collagens, in view of the available genetic reagents and evidence that collagens usually tolerate C-terminal tagging. Collagens make up about half the *C. elegans* cuticle by mass.^57^ Less is known about non-collagenous proteins such as cuticlins or epicuticlins, components of the outer cortical layers.^21,78^ Tagging of Hedgehog-related proteins has also revealed specific localization to substructures of the cuticle or precuticle.^79^ Using collagen localization as a reference, tagging of additional non-collagen aECM components such as ZP domain cuticlins or lectins should be highly informative.

### Optimization of CRISPR for large scale knock-in projects

Our aECM toolkit establishes numerous lessons to establish large-scale tagging projects. Our fastest screening approach involved RNPs, PCR-derived linear repair templates, and screening under a fluorescent dissecting scope. We were most successful with genes expressed at high levels (>1000 transcripts per million in a single-cell RNA-seq data set^2^) and in large tissues such as the epidermis. Interestingly, neither *dpy-10* co-conversion nor a dominant *rol-6(su1006)* co-injection marker helped enrich for aECM knock-ins, perhaps because such cuticle mutants display genetic interactions with aECM target genes. KIs with spatially or temporally restricted expression were much easier to identify with our selection-based approach.

Approximately 10% of the genetically defined cuticle collagen strains displayed overt or cryptic phenotypes, suggesting C-terminal tagging of most collagens is mostly tolerated. Some collagens such as SQT-3 undergo C-terminal processing by the protease DPY-31,^62^. Although we avoided tagging SQT-3 at the C-terminus, other collagens with predicted C-terminal cleavage sites (see Methods) appeared to tolerate tagging. Some collagens proved challenging to tag due to lack of unique crRNA sites. Our C-terminal QUA-1 knock-in was likely processed and an internal tagging attempt broke QUA-1. While our RNP and *unc-119(+)-*based selection approaches are high-throughput, SEC-based selection^80^ with longer homology arms could be a useful alternate approach to tagging proteins that have highly similar homologs.

### Uses of the resource and prospects for future tagging resources

Our resource provides the first large-scale overview of aECM protein localization. Our resource can address key questions in aECM biology such as how the aECM is re-established in wound healing, how it is maintained during normal adult aging, how proteases and protease inhibitors coordinate its remodeling, and how specific aECMs are assembled in a cell type specific manner.^71^ Importantly, pathogenic variants^81^ can now be modeled using proteins expressed at physiological levels.

The *C. elegans* matrisome includes ∼700 members of conserved matrix families,^9^ including receptors. Other potential matrix components are not highly conserved in primary sequence but are conserved in terms of intrinsic disorder propensity or low sequence complexity. For example the *C. elegans* pharyngeal cuticle contains numerous intrinsically disordered proteins,^82^ and epicuticlins consist solely of low complexity repeats.^21,83^ Taking 700 as a lower bound of protein species in the *C. elegans* ECM, at most 25% have been tagged in this work and other projects. The matrisome itself is a small fraction of the ‘secretome’ defined as all proteins normally secreted into extracellular compartments, ∼3500 proteins in humans.^84^ The distinction between matrisome and secretome proteins is not rigid, and proteins may form a continuum of stability or diffusibility within extracellular space. Localization of most secretome proteins has not been investigated using protein tagging. Such approaches could be highly informative for understanding the architecture of the extracellular environment in development, adult homeostasis, wound healing, and aging.

### Limitations of the study

Our toolkit is based on tagging endogenous loci with fluorescent protein tags, and thus conclusions are based on the localization of the tag itself rather than the endogenous protein. We detected an effect of tagging on gene function in a small minority of cases, however it is possible that many more tags have subtle effects on function not detectable in the single mutants. Fluorescent tags could alter stability or localization of the tagged protein. In most cases we used a single tag site and have not validated multiple sites. We have validated numerous knock-ins at the site of insertions but have not searched exhaustively for aberrant editing. Off-target effects of CRISPR/Cas9 editing in *C. elegans* are very rare, but complex alterations at the editing site are sometimes seen^85^. In any event, conclusions based on a single knock-in strain should be treated with caution.

## RESOURCE AVAILABILITY

### Lead contact

Requests for further information and resources should be directed to and will be fulfilled by the lead contact, Jordan Ward

### Materials Availability

Knock-in strains reported here are homozygous viable and have been deposited at the CGC, along with information on the insertion site, fluorescence visibility, and other phenotypic information. Plasmids for the modular knock-ins and color swaps will be made available through AddGene. Repair template and knock-in sequences are available upon request. Strain genotypes, plasmid and oligonucleotide sequences, and details of new alleles used in this study are in **Supplemental Tables**.

### Data and Code Availability

Primary data will be available in a standard repository.

## STAR METHODS

### KEY RESOURCES TABLE

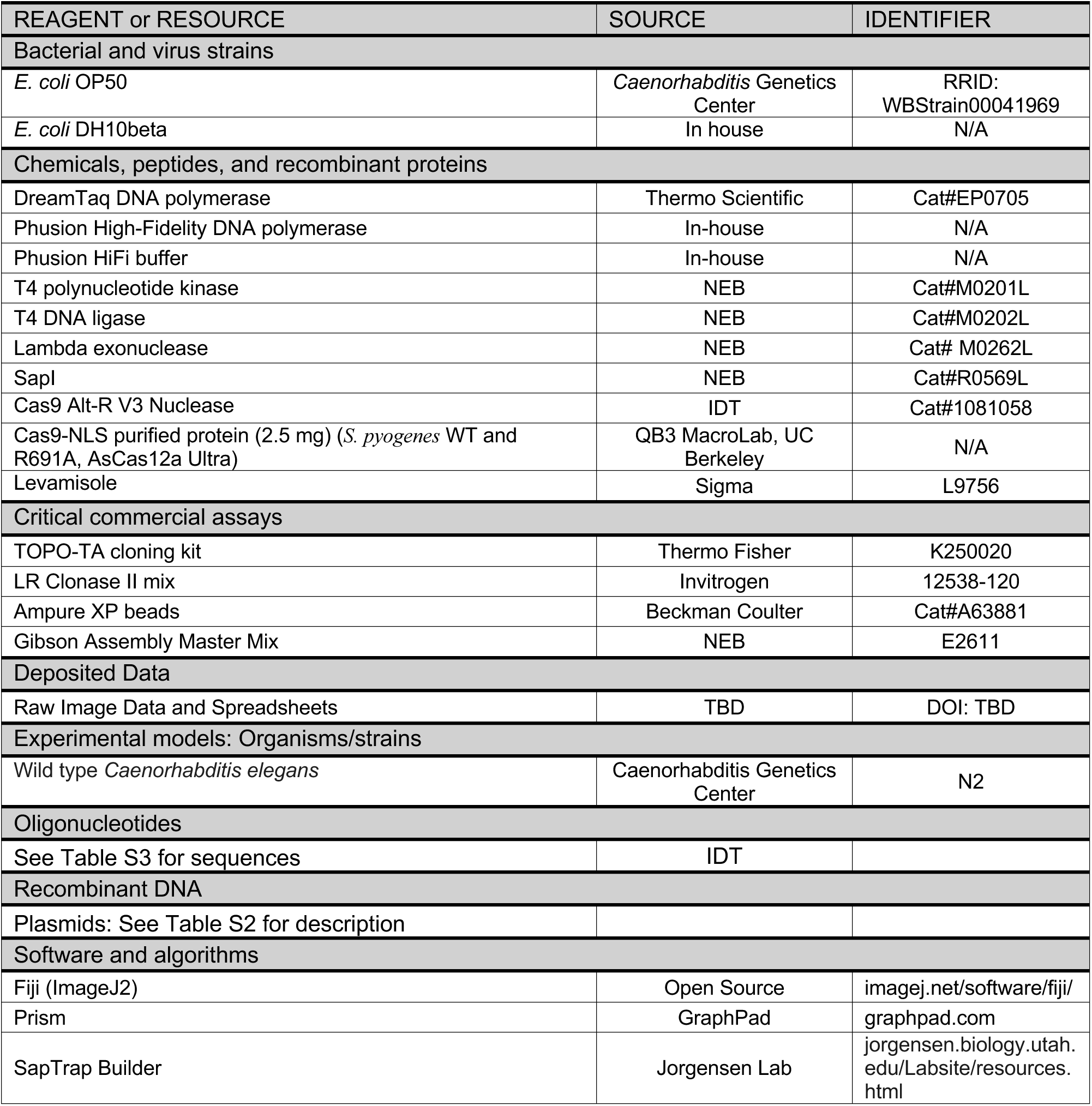

### EXPERIMENTAL MODEL

*C. elegans* were maintained at 20°C on NGM plates seeded with *E. coli* OP50, using standard procedures unless stated. Mutations were confirmed by PCR or sequencing. Strains used in this study are listed in Table S1.

### METHOD DETAILS

#### Molecular biology

We designed flexible 36 bp 5’ and 3’ homology arms encoding N-SSSSSSGGSGGSA-C and N-AASSGSSSGGSGG-C, respectively. These arms were added to the mNeonGreen::3xFLAG from pJW2172 ^86^ by PCR and the products were Gibson cloned into pDONR221 to generate pJW2332. We also designed an mNeonGreen::AID*::3xFLAG equivalent for tagging proteins with nuclear or cytoplasmic domains using a pJW2171 template to generate pJW2333. These cassettes include 4 unique crRNA sites to allow re-editing. pJW2332 was linearized by PCR introducing silent mutations in the PAM to disable the internal crRNA sites. We then cloned in SNAPTag and HaloTag cassettes to make pJW2335 and pJW2336, respectively. We also designed a set of mScarlet cassettes to replace the modular mNeonGreen knock-ins. First, we replaced the 3xMyc cassette from pJW2072 (30 amino acid linker:mScarlet(GLO)::3xMyc::10 amino acid linker)^86^ with 3xHA and 3xOLLAS tags through Gibson cloning to make pJW2469 and pJW2470, respectively. We then linearized pJW2335 and cloned in the mScarlet(GLO)::3xHA to make pJW2471. We attempted the same with an mScarlet(GLO)::3xOLLAS cassette but only were able to obtain a plasmid with 2xOLLAS (pJW2558). To make a SapTrap compatible modular mNG cassette, we first Gibson cloned the modular mNG::3xFLAG cassette with the 36 bp homology arms into pDONR221 with TCC and GGT connectors for SapTrap cloning to make pJW2474. We then linearized this plasmid by PCR and used Gibson cloning to insert a syntron+ *C. briggsae unc-119(+)* cassette from pMLS271^41^ such that the cassette is in an antisense orientation to the mNG cassette, yielding pJW2476.

dsDNA repair templates with 35 bp homology arms were generated for RNP-based genome editing by PCR amplification. 8-12 50 µl PCR reactions were performed and pooled then purified using Ampure XP beads as previously described.^87^ ssDNA repair templates with 35 nt homology arms were generated by PCR amplification similar to the dsDNA repair templates except that one primer was phosphorylated on the 5’ end. This phosphorylation was either performed during primer synthesis (Invitrogen) or in-house using T4 polynucleotide kinase (NEB). The phosphorylated strand was degraded using lambda exonuclease (NEB) and purified as previously described.^37^ For some edits, ssDNA or 5’ biotinylated dsDNA repair templates drastically improved editing efficiency. Our cassette swaps particularly benefitted from using ssDNA or biotinylated DNA templates (**Figure 1**). Notably, we got efficient editing with 35 nt homology arms with our ssDNA templates, whereas a previous report used 59-134 nt homology arms. The shorter homology arms should substantially cut down reagent costs.

*unc-119*-selection based repair templates were generated by SapTrap cloning.^41^ Oligos to generate a 20 bp sgRNA targeting sequence and 60 bp 5’ and 3’ homology arms were designed by SapTrap Builder.^88^ We set the program to produce a C-terminal fusion and removed the “5’-CGC-3’” connector at the 5’ end of the hUS-B oligo with “5’-GGA-3’” to make this homology arm compatible with our modular toolkit. We performed similar SapTrap reactions as described^41,89^ with minor modifications. To anneal the oligonucleotides, we added 1 μl of each 100 µM oligo stock (100 pmol/μl) to 8 μl of water and pipetted up and down to mix. We then added 0.5 μl of that dilution to 5 μl 10x annealing buffer (500 mM KOAc, 50 mM Tris pH 7.4) + 44.5 µl dH_2_O. We then ran on an annealing program on a thermocycler consisting of 95°C for 5 min followed by dropping 1°C per minute down to 4°C. Annealed oligos were stored at 4°C before use. We made 10x SapTrap buffer by adding 2 µl of 2.5 M potassium acetate to 48 µl of 10x T4 ligase buffer. We thawed a fresh tube of 10x T4 ligase buffer and made ∼50 µl aliquots. For both the 10x SapTrap buffer and 10x T4 ligase buffer we tally each freeze-thaw on the tube as part of our quality control pipeline. We made a 10x SapTrap enzyme mix by mixing 10 µl of SapI (pipetting up and down repeatedly before transferring), 5 µl of T4 PNK (NEB) and T4 DNA ligase (NEB). If multiple stocks were present in the lab, they were marked and used oldest to newest. We then set up SapTrap reactions with ∼20 fmol of pDD379^90^ (backbone with F+E sgRNA; ∼50 ng), ∼20 fmol of pJW2476 (modular mNG::3xFLAG+*unc-119(+)*; ∼75 ng) and 200 fmol of each annealed oligo pair (5’ homology arm, 3’ homology arm, sgRNA). While establishing an efficient editing pipeline we set up a control omitting pJW2476 and the annealed oligos. We ran the reactions on a thermocycler as follows: 37°C for 10 min, 16° C for 5 min followed by 30 cycles of 37°C for 5 min, 16°C for 5 min. There was a final 37°C 10 min digestion followed by 65°C for 20 min. To increase cloning efficiency, we added 1 µl of SapI (pipetting up and down before transferring) and incubated at 37°C for two hours followed by a 65°C 20 min heat kill step. The entire reaction was transformed into in-house made DH10beta competent cells and plated on LB+Amp. We frequently screened reactions with colony PCR using M13-Rev (5’-CAGGAAACAGCTATGAC-3’) and 5’-AGCTCCTCGTATCCGTCGTTTG-3’. Colonies were picked into 50 µl of sterile water in a 12-well PCR strip or 96-well plate. 1 µl of this mixture was used as a template in a 10 µl PCR using homemade Phusion PCR and 5x Phusion HiFi buffer with green loading dye. We typically use a pJW2586 *col-75* repair template as a positive control in colony PCR reactions.

#### CRISPR knock-ins

JDW strains were generated by the Ward lab using the following methods. crRNA, trcRNA (both from IDT) and Cas9 (either IDT Alt-R Cas9 V3 Nuclease or UC Berkeley MacroLab) were mixed and incubated at 37 °C for 15 min. Note that a range of protein and RNA were used in the optimization (**Figure 1**). Pre-optimization reactions used 700 ng of IDT Cas9 and a 2:5:2.5:1 ratio of crRNA:trcRNA:Cas9. Post-optimization reactions used 250 ng of IDT Cas9 and a 1:1:1 ratio of crRNA:trcRNA:Cas9. Repair template PCRs were melted as described^87^ and used at 50 ng/µl. ssDNA repair templates were used at 110-250 ng/µl. We tested *dpy-10* co-conversion and pRF4 co-injection markers using previously described approaches.^1,38^ We also developed an alternate co-conversion approach using insertion of a T2A::GFP cassette at the 3’ end of *myo-2* to create an endogenous promoter reporter.^91^ Optimized editing used 10 ng/µl of a pSEM232 (*mlc-1p::2xNLS::tagRFP::tbb-2* 3’UTR) co-injection marker.^92^ Animals were screened for mNeonGreen expression on a Leica M165 FC GFP/RFP stereomicroscope with 16.5:1 zoom ratio, 7.3-120x magnification range and GFP or GFP long pass filters, and an Excelitas X-Cite FIRE LED light source. We generated 10 color-swapped strains as a proof of principle using a biotinylated linear dsDNA repair template and the internal crRNA sites. We chose this repair template as it boosted efficiency and we tended to get more injections out of a single prep compared to ssDNA. PHX strains were generated by SunyBiotech.

*unc-119(+)-*based selection was performed by injecting repair template plasmids containing sgRNA (F+E) cassettes^90^ at 65 ng/µl, 25 ng/µl pCFJ2474^93^ (*smu-1p::Cas9(PATCs)::gpd-2 tagRFP-T(myr, PATCs)::smu-1 3’UTR*), 10 ng/µl pSEM238^94^ (HisCl counter-selection marker), 25 ng/µl pSEM232 (*mlc-1p::2xNLS::tagRFP::tbb-2* 3’UTR)^92^ into CFJ302, a temperature-sensitive *unc-119(kst33)* strain. We made use of a recently generated temperature-sensitive *unc-119* allele which further facilitates adoption of this pipeline as labs can inject into animals with a WT phenotype. *unc-119(kst33)* mutants were maintained at the permissive temperature of 15°C, at which they are phenotypically wild type, prior to injection. After injection, animals were shifted to the non-permissive temperature of 25°C at which a fully penetrant Unc phenotype manifests, unless animals are rescued with *unc-119(+).* Plates were flooded with histamine when they starved and wild-type moving animals lacking the *mlc-1p::tagRFP* marker were picked as candidate knock-ins. The selective marker was excised through injection of 10 ng/µl pMDJ20 (*smu-1p::Cre Recombinase (PATCs):: smu-1 3’ UTR*]),^93^ 10 ng/µl pCFJ782^95^ (HygroR marker), 25 ng/µl pSEM232. Unc animals were picked, outcrossed to remove *unc-119(ts)*, and the knock-in recovered by either fluorescence or PCR. The *spia-1* mNG::3xFLAG knock-in was generated using an SEC-based approach.^44^ Modular mNG::3xFLAG knock-ins for *bli-1* and *noah-1* recapitulated independently generated knock-ins and have been described.^96^

We observed an interesting variation in repair template efficiency that warrants further investigation. With our *glh-1* GFP knock-ins, we did not observe a difference in efficiency with unmodified dsDNA, 5’ modified dsDNA, or ssDNA repair templates (**Figure S1C,D**). In contrast, for *glh-1* mNeonGreen::3xFLAG to mScarlet::2xOLLAS color swaps the biotinylated dsDNA repair template was 10x more efficient and the ssDNA repair template 34x more efficient than an unmodified dsDNA template, respectively (**Figure 2**). As *glh-1* is an efficient edit, modification might not further improve efficiency. Notably, unmodified dsDNA repair templates allowed efficient tag swapping with other combinations of crRNAs (**Figure 2**). For large-scale tagging we used unmodified dsDNA repair templates as they were cheaper than biotinylated repair templates and faster to produce than ssDNA repair templates. However, for smaller scale editing or edits that prove challenging such alternative templates may be a better approach.

#### Sequence validation

All knock-ins were validated by PCR using primers that bind outside of the homology arms followed by Sanger sequencing. For knock-ins with *unc-119(+)* selection we initially performed PCRs to amplify the 5’ and 3’ knock-in junctions and Sanger sequenced the PCR products. Following selectable marker excision, we PCR amplified the insertion and performed Sanger sequencing.

#### Collagen domain organization, sequence classification, and cleavage sites

Cuticle collagen genes are defined as encoding proteins with a central block of 37-43 Gly-X-Y triplet repeats (with one or more short interruptions) and additional motifs such as a dibasic subtilisin cleavage site (Homology box A), three Cys-rich clusters, and a signal peptide or transmembrane domain. Approximately 35 collagens have potential C-terminal cleavage sites resembling that of SQT-3 (DGG motif) including BLI-6, DPY-4, DPY-13, ROL-1, and LON-3. ROL-8 has been classified as non-cuticular^9^ however it has several hallmarks of cuticle collagens including an N-terminal dibasic cleavage site. COL-135 has an atypically large region of ∼200 triplets, most of which are Gly-X-Lys.^9^ COL-55 has a short Gly-X-Y region and is not represented in RNAseq data; it contains a dibasic cleavage site and a Cys-rich motif. COL-70 lacks Gly-X-Y repeats but instead contains a repeated motif GGPG(N/T)P. We excluded *col-182* as it is a pseudogene in the domesticated N2 strain.^97^ We excluded known non-cuticular collagens such as LET-2, EMB-9, CLE-1, MEC-5, and transmembrane proteins such as COL-99, UNC-122, and COF-2.

#### Whole Genome Sequencing, Sanger Sequencing, and editing of *col-71*

We performed whole-genome sequencing (WGS) of the BE22 strain, containing the roller allele *sc22*. Genomic DNA was prepared as described^98^ and resequencing performed at BGI Americas (San Jose, CA). We inspected BE22 sequence specific variants for genic changes on chromosome II relative to N2 wild type. Sanger sequencing of alleles of genetically defined collagens used standard procedures. To edit the *sc22* missense change in *col-71* we used the melting method.^87^ We injected N2 animals with 2 µM *col-71* crRNA and 4 µM of repair template along with 50 ng/µl P*inx-6*-RFP co-injection marker and Cas9 (UC Berkeley MacroLab). F_1_ progeny were picked based on the co-injection marker and the *ju2038* SNP mutation verified by sequencing. Positive edits were retrieved as heterozygotes; we picked Rol progeny and verified the mutation again by sequencing.

#### Imaging

We assessed overall strain brightness and localization using Leica MZ FLIII or M165 fluorescence dissection microscopes. For widefield and confocal imaging we immobilized animals using 10 mM levamisole and imaged them on a Zeiss Axio Imager M2, a Zeiss LSM800 confocal, or a Zeiss LSM900 confocal with Airyscan capability, as described.^21^ We varied laser power depending on the brightness of the specific knock-in strain. For the brightest strains we reduced intensity by cutting the emission window. Confocal z stacks were generally made from 5-7 planes 0.3 µm apart to image the entire adult cuticle thickness. For imaging of the fibrous layers we used z sections 0.17 µm apart. Airyscan speeds varied from 7 (fast) to 5 (slow); samples were first imaged using the fast setting, then more slowly for higher resolution as needed. Unless stated, animals were staged in mid L4 based on vulval morphology then imaged 24 h later; images in Figures are representative of at least 10 animals imaged per strain. FRAP experiments were performed as described.^21^

Localization in L4 or adult males was generally analyzed in mNG/+ heterozygous cross progeny from crosses of N2 males with mNG/mNG hermaphrodites. For quantitation of knock-in expression intensity in knock-in heterozygous animals, we first crossed N2 wild type males with mNG/mNG homozygotes, then crossed mNG/+ heterozygous F_1_ males with N2 hermaphrodites. Fluorescence intensity was measured in at least 3 ROIs in the midbody lateral cuticle of F_2_ animals 24 h post mid L4 stage. Background was subtracted and data normalized such that the homozygous strain mean = 1.

#### Quantitation and statistical analysis

Statistical analyses were performed using GraphPad Prism 10. Data were tested for normality, and parametric or non-parametric tests used as appropriate. Statistical significance was determined using Fisher’s exact test, unpaired t test or Mann-Whitney test. For multiple comparisons we used one-way ANOVA or Kruskal-Wallis tests followed by a post test. Data are represented as mean ± SEM unless noted. Sample sizes are shown inside or above columns in Figures or legends.

## ACKNOWLEDGMENTS

We thank members of our labs and David Sherwood, Helge Grosshans, Yishi Jin, Andreas Ernst, Nathalie Pujol, Cathy Savage-Dunn, King Chow, Max Heiman, Emily Troemel, and Meera Sundaram for discussions or comments on the manuscript. We thank Jennifer Adams, Jade Diaz, Risa Iwazaki, Erin Jyo, Vanessa Lambatan, Xinran Shi, and Charlotte Sue for additional strain construction and characterization, Laura Toy for sequencing, and Patricia Bliatout for lab support. Funding: NIH R35GM134970 (ADC); R21OD033663 (JDW and ADC); R01GM138701 (JDW); R35GM158317 (JDW).

## AUTHOR CONTRIBUTIONS

J.D.W. and A.D.C. conceived the project. M.P. performed strain construction and confocal imaging. K.K. and C.C. performed strain construction, quantitation, and imaging. J.M.R. performed microinjections to create all JDW strains. J.M.R., G.E.A, E.C., B.B., A.R.B., S.H.M, and T.E.W. created repair templates and validated JDW knock-in strains. J.D.W. and A.D.C. secured funding. All authors reviewed and edited the paper.

## SUPPLEMENTAL INFORMATION

### Supplemental Tables

**Table S1.** Strains, new alleles, and genotypes

**Table S2.** Plasmids

**Table S3.** Oligonucleotides

**Table S4.** CRISPR reagents

### Supplemental Figures

**Figure S1.**
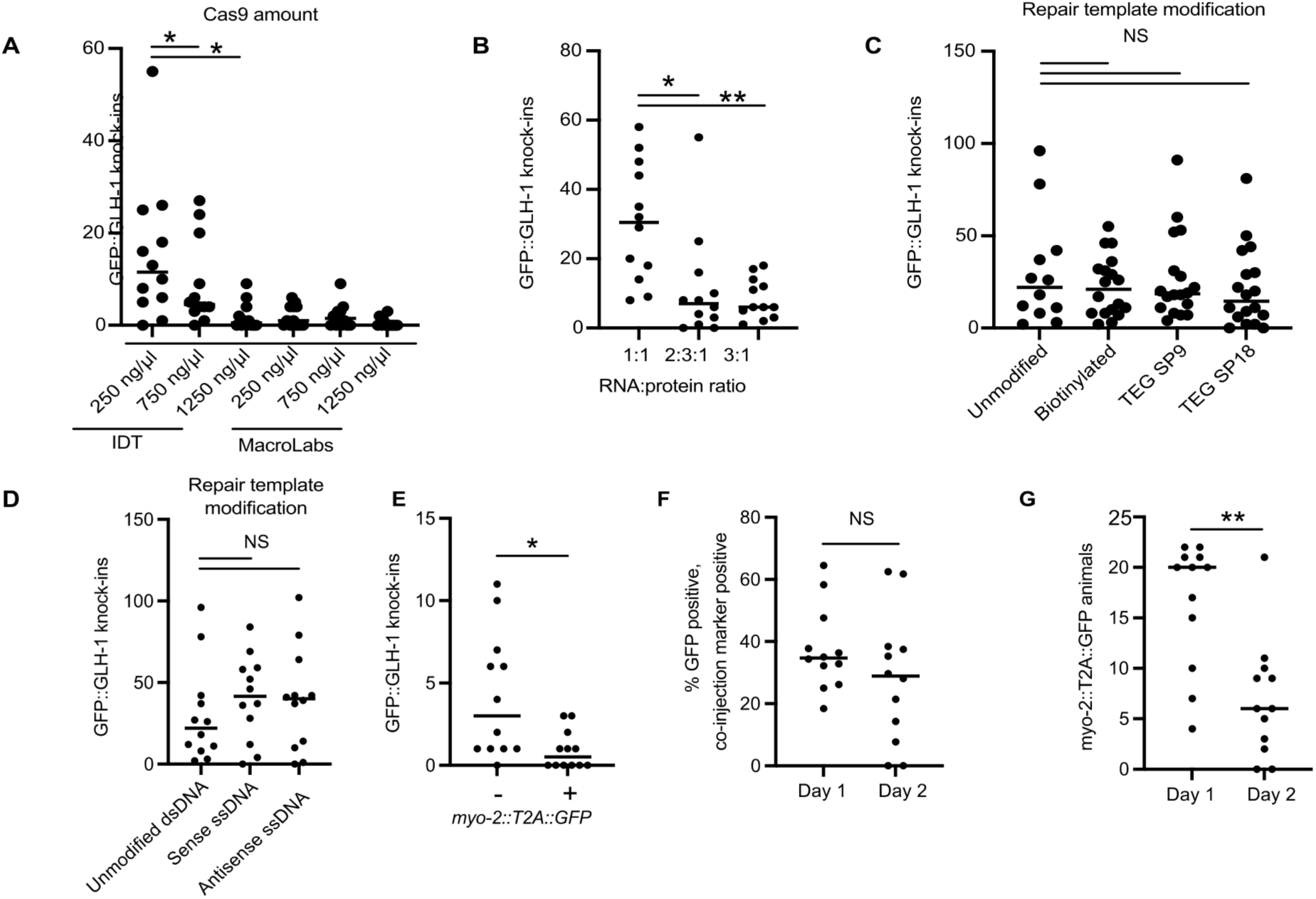
Optimization of knock-in editing parameters. (A-D) Editing optimization using GFP::GLH-1 knock-ins as a test case. (A) Cas9 from the indicated sources at the indicated amounts with a 2.5:1 RNA:protein ratio. IDT Cas9 at 250 ng/µl was most efficient, similar to previous reports.^1^ (B) The indicated RNA:protein molar ratios (crRNA and trcRNA at equal concentration, RNA amount varied to 250 ng/µl Cas9). (C and D) The indicated repair templates with 250 ng/µl Cas9, 1:1 RNA:protein ratio). (E) Number of GFP::GLH-1 knock-ins found in animals lacking (-) or containing (+) a *myo-2::T2A::GFP* co-CRISPR knock-in. (F) Percent of *mlc-1p::tagRFP* co-injection marker positive animals laid on day and day 2 post-injection with *myo-2::T2A::GFP* knock-ins. (G) Number of *mlc-1p::tagRFP* co-injection marker positive animals laid on day and day 2 post-injection with *myo-2::T2A::GFP* knock-ins. Statistical significance was determined using a two-tailed unpaired Student’s t-test. *P* < 0.05 was considered statistically significant. * Indicates *P* < 0.05, ** indicates *P* < 0.005. Two independent experiments were performed for all optimization tests.

**Figure S2.**
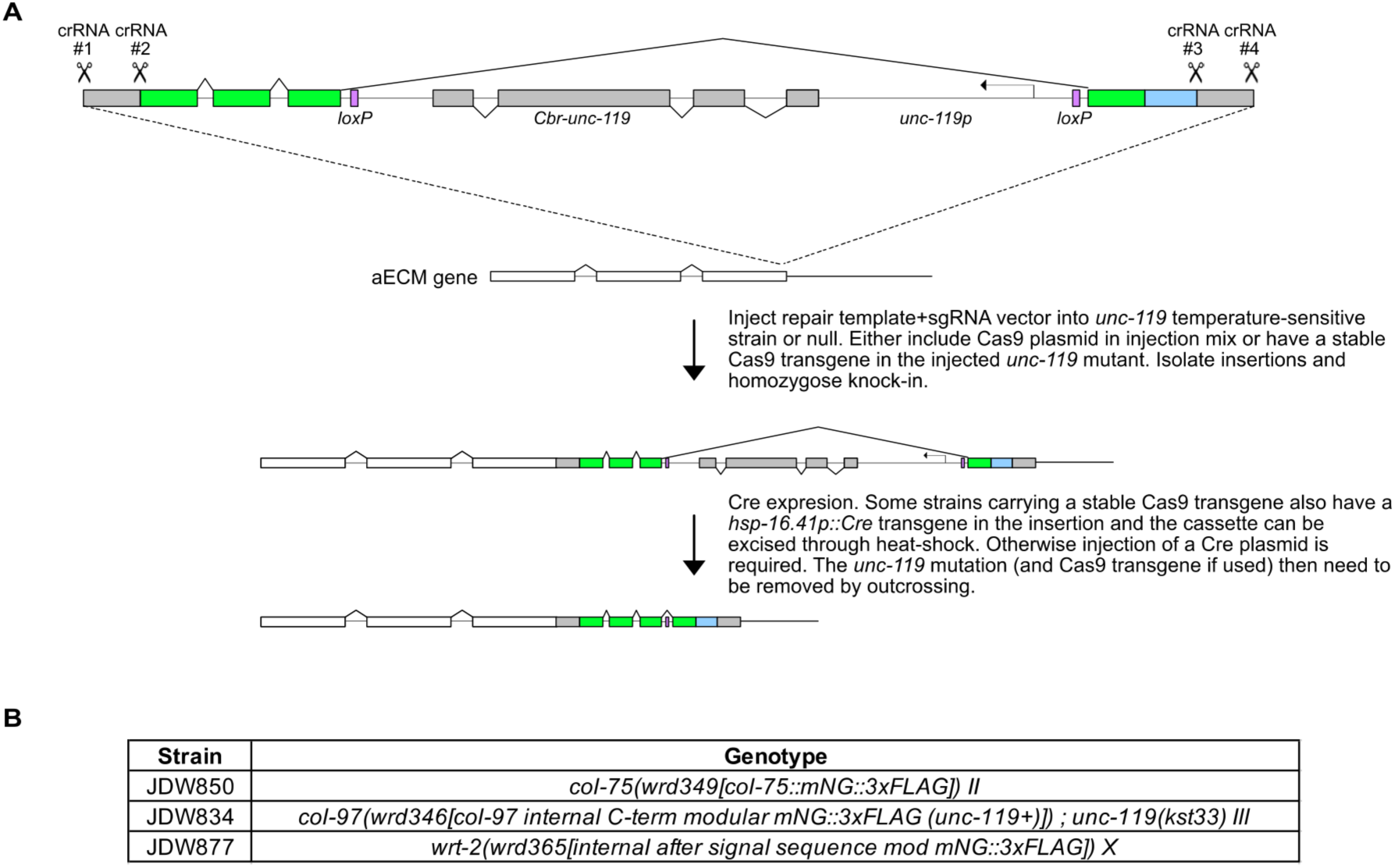
An *unc-119(+)-*selection based modular mNeonGreen::3xFLAG knock-in pipeline. (A) A schematic of the knock-in cassette with the *C. briggsae unc-119(+)* rescue cassette being transcribed from the opposite strand as the mNeonGreen::3xFLAG cassette. The selection cassette is flanked with loxP sites and is within an intron. Knock-ins rescue the uncoordinated phenotype of *unc-119* mutants and the cassette can be excised through Cre expression. (B) Strains generated with the *unc-119(+)* knock-in pipeline. Our RNP-based pipeline was most efficient with genes expressed in larger tissues (hyp7 syncytium, seam cells, pharyngeal cells) and at high levels (>1000 transcripts per million in a single cell L2 stage RNA-seq dataset^2^ as these criteria allowed facile screening under a fluorescence dissecting microscope. *unc-119(+)* selection is more suitable for high throughput pipelines than alternatives such as self-excising cassettes, as the 60 bp homology arms can be made using oligo annealing, bypassing need for PCR. We developed a robust assembly pipeline and successfully tagged two collagens (*col-75* and *col-97*) and one warthog gene (*wrt-2)* as proof-of-principle. We chose these test cases as: i) *col-75* is predicted to be restricted to socket cells and excretory duct or pore cells at moderate levels; ii) an RNP-based attempt at *col-97* tagging had failed; and iii) *wrt-2* was expressed at moderate levels.

**Figure S3.**
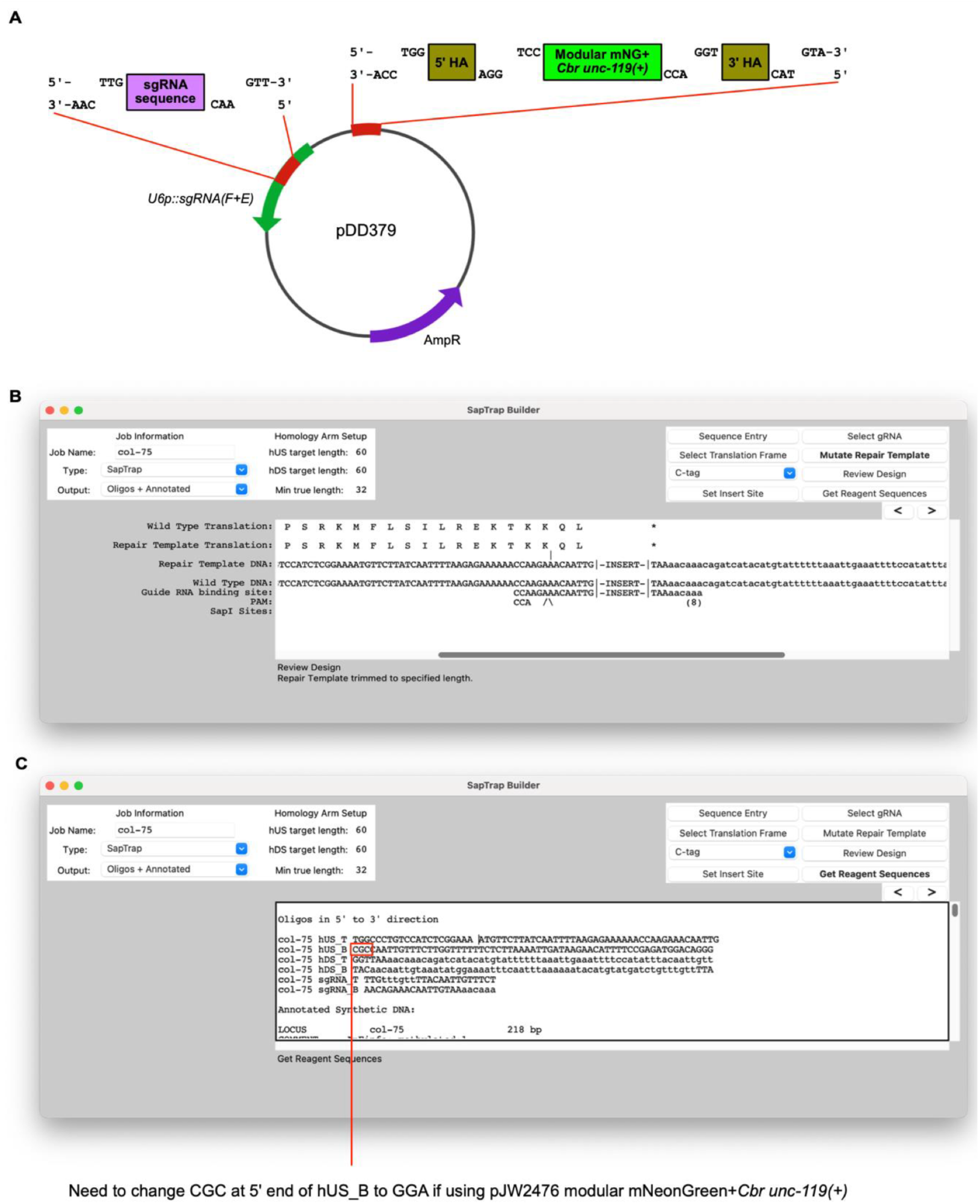
A SapTrap assembly pipeline for *unc-119(+)-*selection based modular mNeonGreen::3xFLAG repair templates. (A) Oligos encoding the target sgRNA, the 5’ and 3’ homology arms (HA) are annealed to produce the indicated 3 bp overhangs and combined with the pJW2476 vector containing the modular mNeonGreen::3xFLAG *unc-119(+)* cassette and an appropriate backbone (such as pDD379) in a SapTrap cloning reaction. (B, C) Using SapTrap Builder to design knock-in reagents. (B) depicts the settings used in SapTrap Builder (60 nt HA oligos, aka hUS, hDS. C- tag), for internal or N-terminal tagging. The C-tag produces the desired 3 nt overhangs. (C) indicates the change to the output needed for a custom 3 nt overhang for one connection, replacing the 5’-CGC-3’ at the 5’ end of hUS_B with “5’-GGA-3’”.

**Figure S4.**
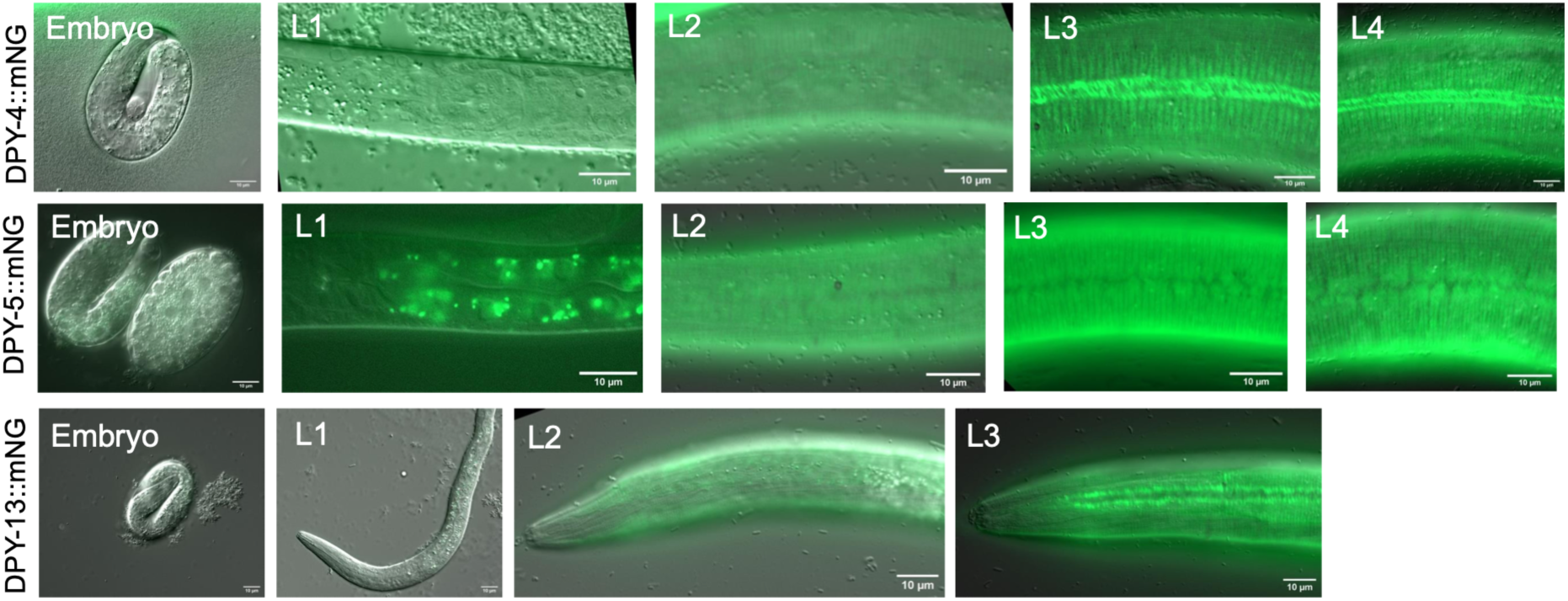
Stage specificity of collagen knock-in expression. (A) DPY-4::mNG, (B) DPY-5::mNG, and (C) DPY-13::mNG are not detectable in late embryos; fluorescence is first detectable in late L1 and remains visible through L4 stage. Widefield DIC and fluorescence. Scales, 10 µm

**Figure S5.**
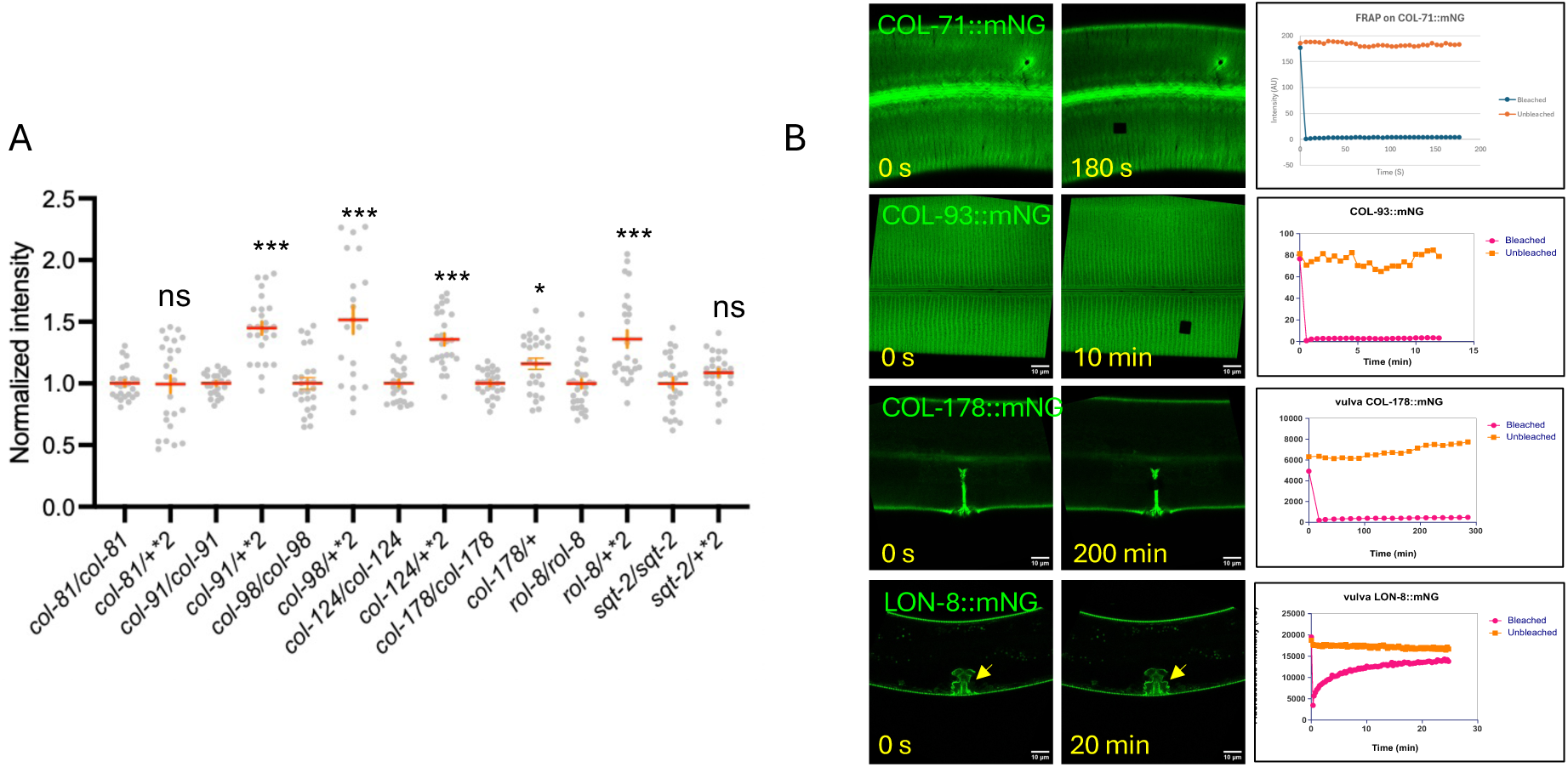
Knock-in dosage and mobility of fusion proteins. (A) mNG fluorescence intensities in the midbody cuticle from homozygous and heterozygous strains, dot plots of intensities normalized to controls (see Methods). Intensities of heterozygous strains were doubled to correct for copy number and compared to the homozygous strain intensity by Mann-Whitney test; ***, P<0.001. (B) Fluorescence Recovery After Photobleaching (FRAP) of selected KIs in the body and vulval cuticles. Most mNG fusion proteins did not show significant recovery, apart from LON-8::mNG in the vulval cuticle.

**Figure S6.**
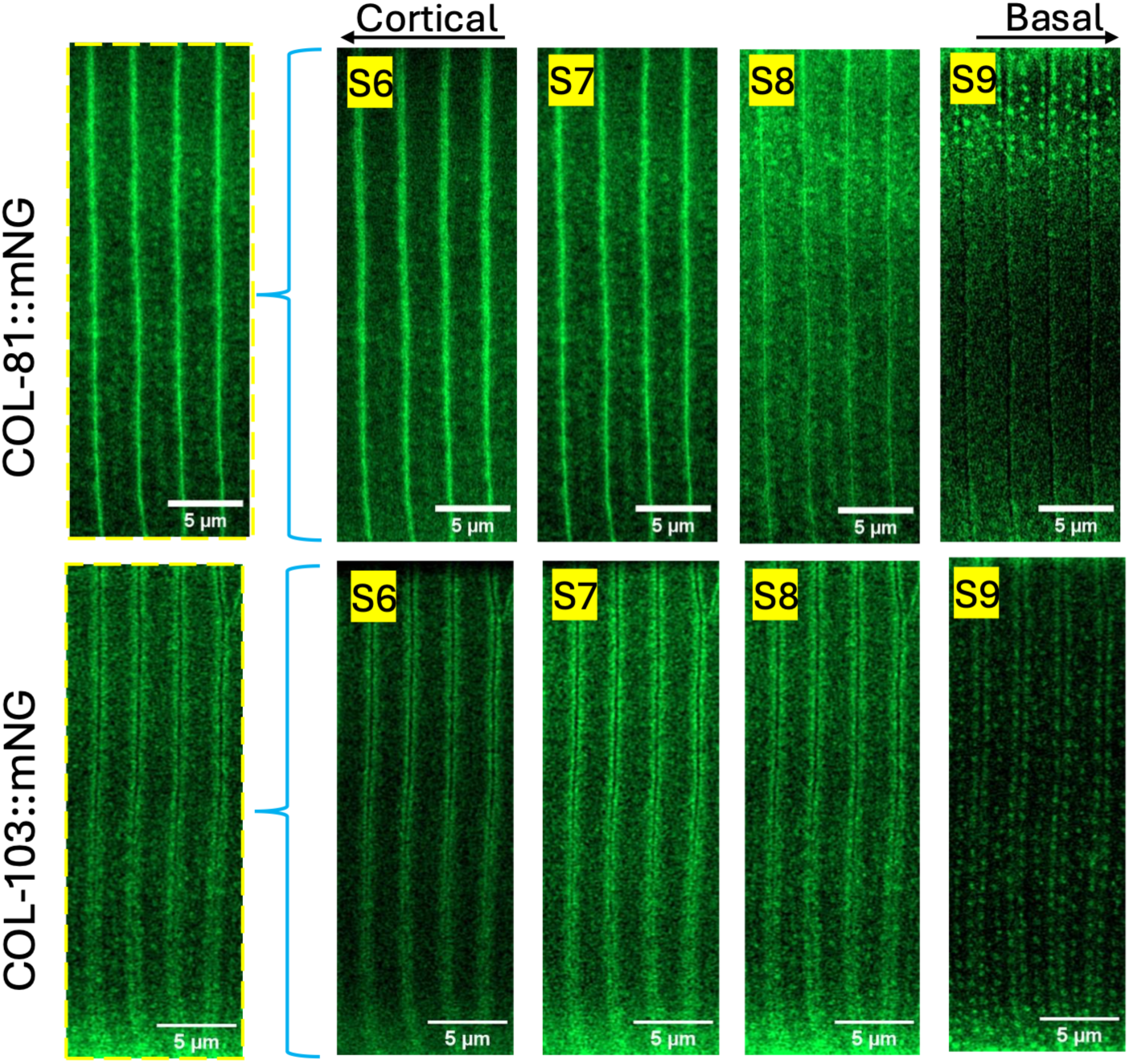
Furrow and strut localization. Single Airyscan focal planes showing dual localization to furrow and struts. COL-81::mNG localizes to the midline of each furrow in cortical planes (S6) and to strut puncta in basal planes (S9). COL-103::mNG localizes to furrow-flanking stripes in cortical planes (S6-8) and to strut-like puncta in basal planes (S9). Scales, 5 µm.

**Figure S7.**
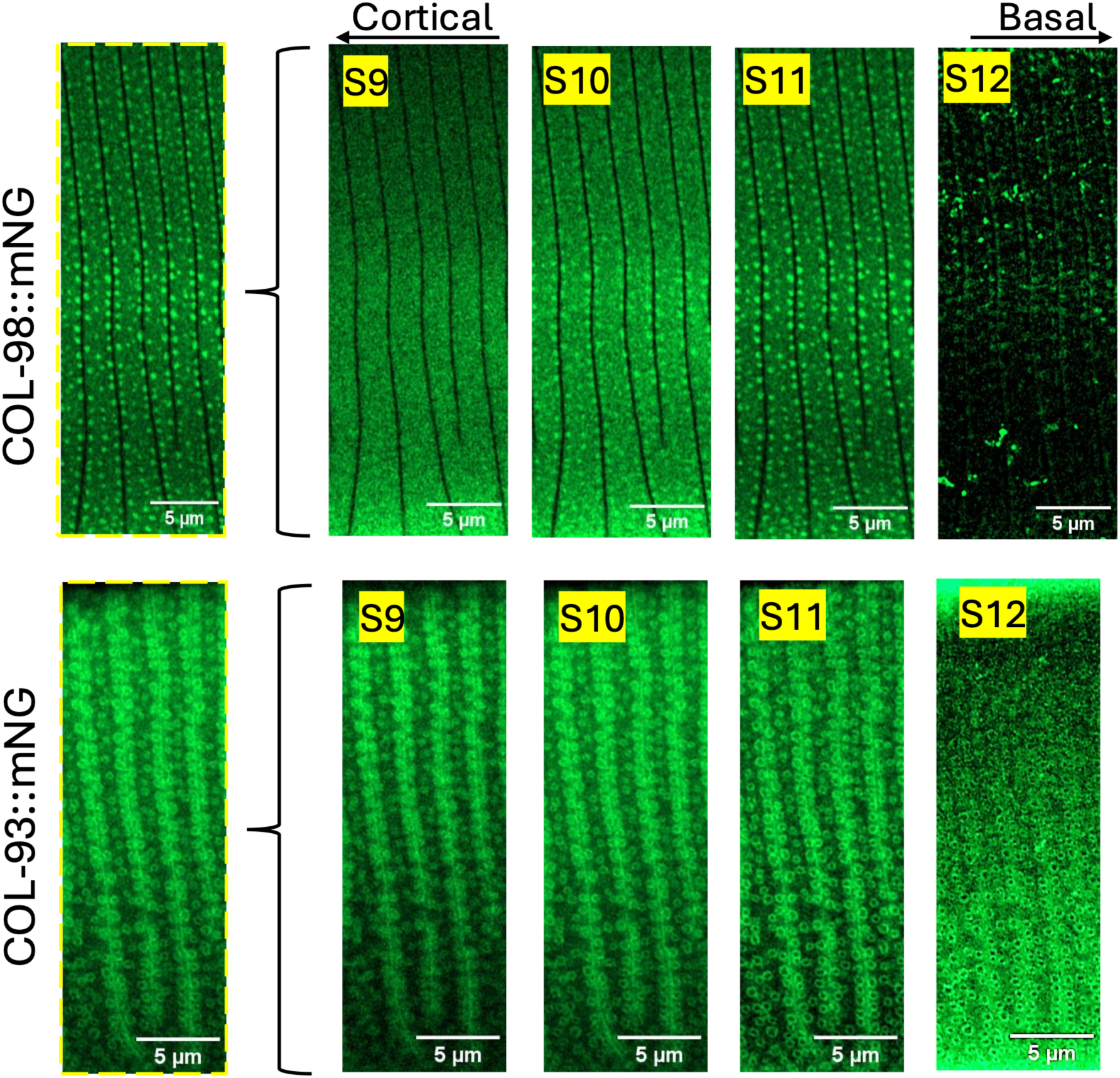
Strut localization. Single Airyscan focal planes of strut-related localizations. COL-98::mNG shows diffuse localization in more cortical planes (S9-10) and punctate strut localization in medial planes (S11). COL-93::mNG shows ring-like localization in multiple focal planes. Scales, 5 µm.

**Figure S8.**
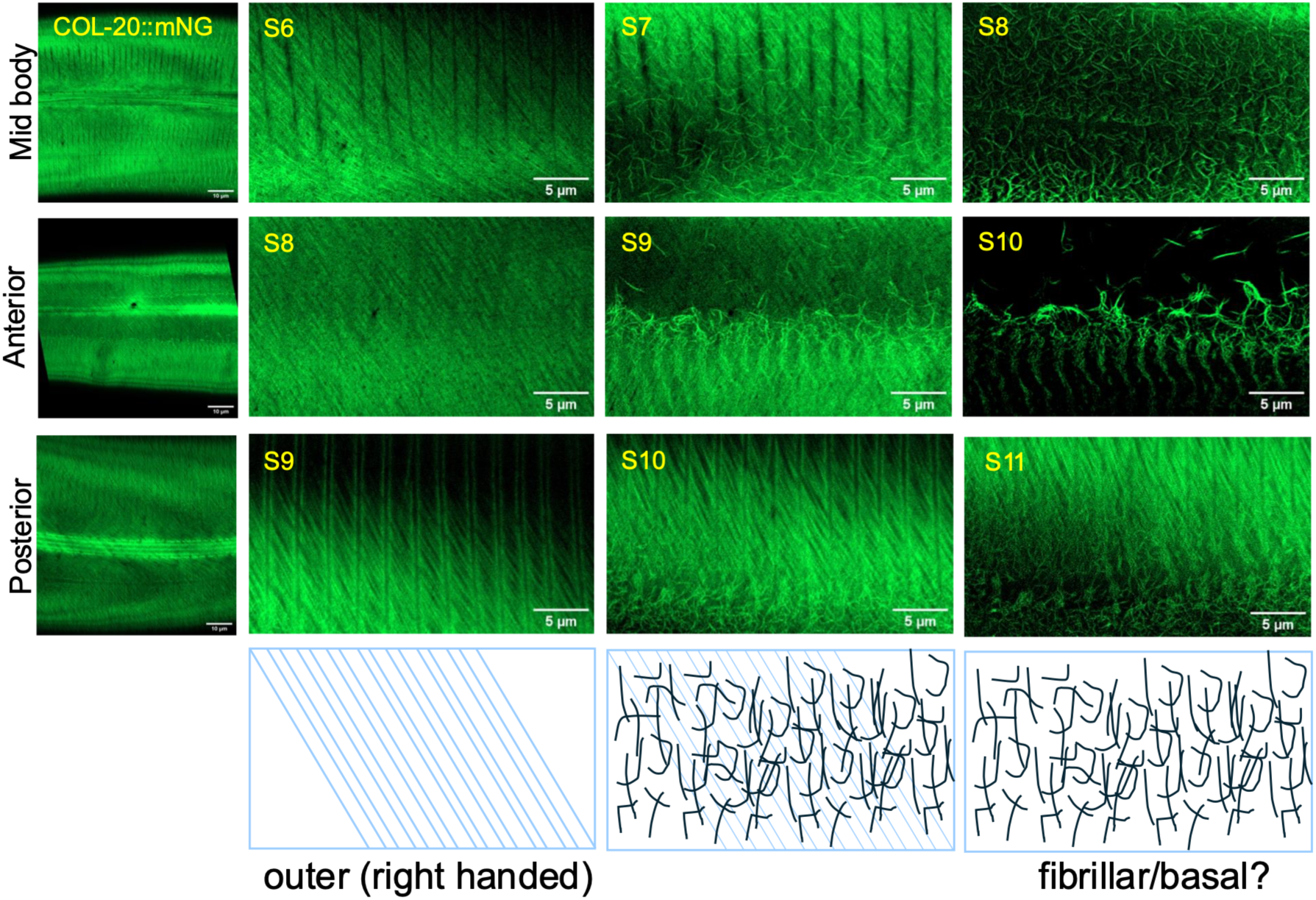
Fibrous layer localization of COL-20::mNG. Airyscan z-stacks and single focal planes of COL-20::mNG showing localization to outer (right-handed) oriented fibrous layer and basal fibrillar layer. Stacks from anterior, midbody, and posterior. Scales, 5 µm. Cartoons of COL-20::mNG localization in different focal planes.

**Figure S9:**
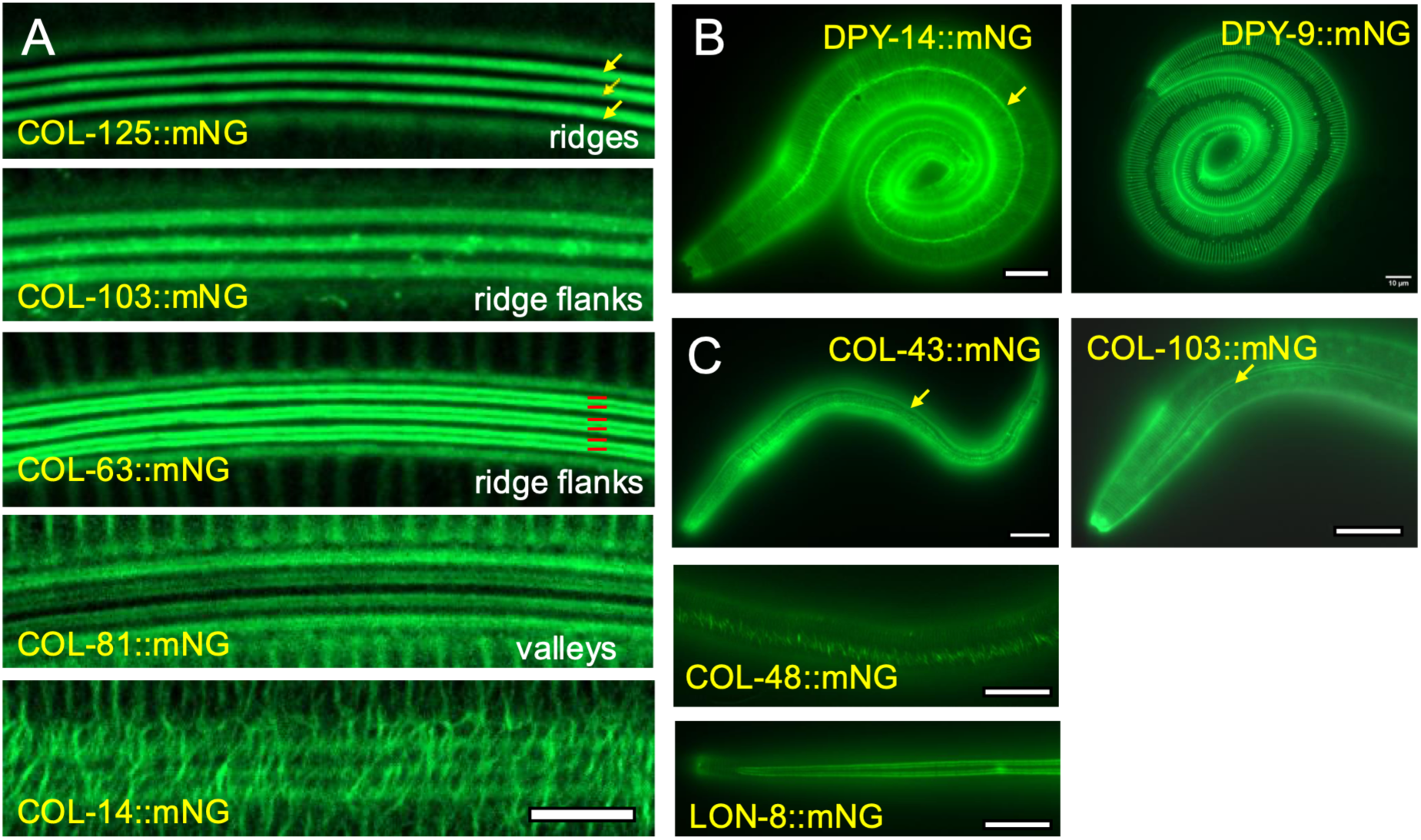
Localization patterns in lateral alae and dauer stages. (A) Selected examples of adult alae localization patterns, showing examples of localization to alae ridges (yellow arrows), ridge flanks (red lines), and valleys. Airyscan projections; scale, 5 µm. (B) L1 alae localization of DPY-14::mNG (yellow arrow) compared to exclusion from alae of DPY-9::mNG. (C) Dauer alae localization of COL-43::mNG, COL-103::mNG, COL-48::mNG, and LON-8::mNG. Scales, 10 µm.

**Figure S10:**
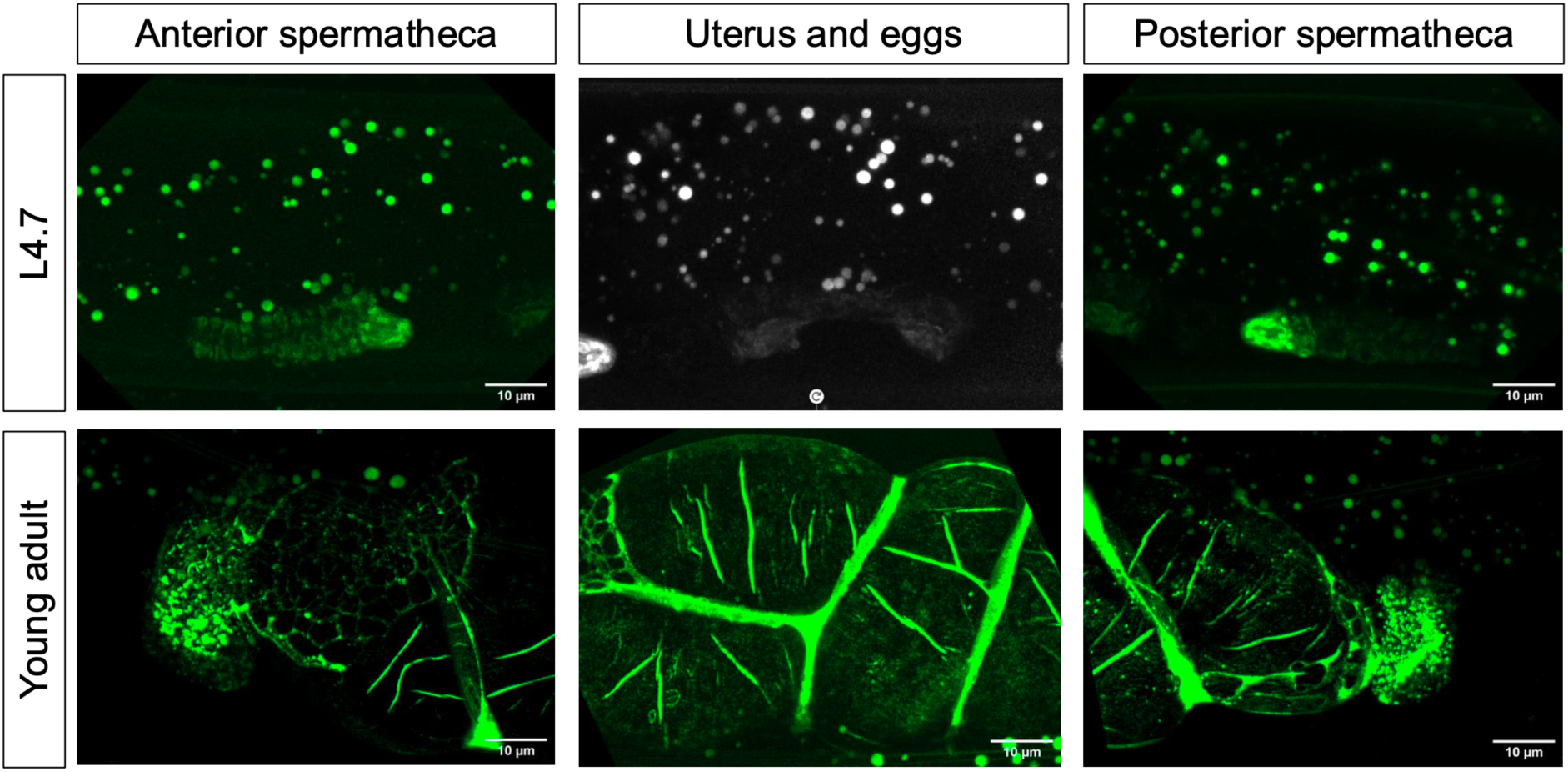
Localization of the atypical collagen COL-135:mNG. Images of anterior spermatheca (left), uterus/eggs (middle), and posterior spermatheca (left) in L4 and young adult stages. COL-135::mNG was expressed in spermathecal cells beginning in late L4 stage (L4.7 shown). In young adults COL-135 localized to uterine matrix surrounding newly fertilized eggs. A small amount of COL-135::mNG appeared to be retained on the eggshell after egg laying. Images are maximum intensity projections of z stacks of 23-49 x 0.5 µm slices. Scales, 10 µm.

